# Comprehensive proteomics analyses identify PIM kinases as key regulators of IL-15 driven activation of intestinal intraepithelial lymphocytes

**DOI:** 10.1101/2020.03.27.011338

**Authors:** Olivia J. James, Maud Vanderyken, Julia M. Marchingo, Francois Singh, Andrew G. Love, Mahima Swamy

## Abstract

Intestinal intraepithelial lymphocytes (IEL) are an abundant population of tissue-resident T cells that protect the gut from pathogens and maintain intestinal homeostasis. The cytokine IL-15 is transpresented by epithelial cells to IEL in complex with the IL-15 receptor α chain (IL-15Rα). It plays essential roles both in maintaining IEL homeostasis, and in inducing IEL activation in response to epithelial stress. IL-15 overexpression also drives the gluten-induced enteropathy Coeliac disease, through cytotoxic activation of IEL. In order to better understand how IL-15 directly regulates both homeostatic and inflammatory functions of IEL, we set up quantitative proteomics of IL-15/Rα stimulated IEL. We reveal that high IL-15/Rα stimulation licenses cell cycle activation, upregulates the biosynthetic machinery in IEL, increases mitochondrial respiratory capacity and induces expression of cell surface immune receptors and adhesion proteins that potentially drive IEL activation. We find that high IL-15/Rα selectively upregulated the Ser/Thr kinases PIM1 and PIM2 and demonstrate that PIM1/2 are essential for IEL to proliferate, grow, and upregulate Granzyme B in response to high IL-15. Significantly, IEL from Coeliac disease patients express high levels of PIM kinases. These unexpected findings reveal PIM kinases to be key determinants of IEL responses to elevated levels of IL-15.

## INTRODUCTION

Intraepithelial lymphocytes (IEL) are a specialised lymphoid compartment in the intestinal epithelium, comprising an extremely heterogeneous population of T lymphocytes. As T cells, IEL express a T cell antigen receptor (TCR) consisting of either αβ or γδ chains, alongside TCR coreceptors CD8αβ or CD8αα and to a lesser extent CD4^(+/-)^. The most prevalent IEL subsets within the epithelium of the murine small intestine are those expressing TCRγδ and CD8αα (TCRγδ CD8αα), which account for ∼50% of the total IEL pool, with the remaining being TCRαβ CD8αβ or TCRαβ CD8αα expressing cells. Residing at the forefront of the intestinal lumen, IEL are exposed to a range of commensal bacteria, dietary antigens and potential pathogens. These immune cells are therefore faced with the conflicting task of protecting the intestinal barrier but also preventing indiscriminate tissue damage. Owing to their tissue-specific roles, IEL are remarkably distinct from conventional CD8^+^ T cells in the periphery. For example, in the resting state IEL have an ‘activated’ T cell phenotype and express high levels of cell surface markers typically only displayed on tissue resident cytotoxic T cells, such as CD44 and CD69 (Shires, Theodoridis and Hayday, 2001). IEL also harbour extremely high levels of cytotoxic molecules such as Granzyme A and B (GzmA and GzmB) under normal conditions, further distancing them from conventional CD8 T cells which only express these proteins following TCR activation. Despite their poised state, IEL do not display effector function under normal conditions and are tightly regulated to maintain their protective role without causing unnecessary damage to the gut epithelium. IEL do not solely depend of TCR stimulation for their activation, rather, signals from the microenvironment are important for communicating compromised barrier integrity to the surrounding IEL and consequently eliciting an effective immune response. One such signal is the common γ-chain (γ_c_) cytokine IL-15.

IL-15 is produced by a wide range of cells including non-hematopoietic cells such as intestinal epithelial cells (Jabri *et al*., 2000; Setty *et al*., 2016) and its expression is elevated in the gut microenvironment during tissue stress or infection (Jabri and Abadie, 2015). IL-15 is most commonly presented to surrounding IEL in a cell contact-dependent manner, known as transpresentation, by epithelial cells expressing IL-15 bound to the high-affinity IL-15 receptor α subunit (IL-15Rα) (Stonier and Schluns, 2010). The signalling components of the IL-15 receptor are made up of the β subunit (CD122/IL-2Rβ), which is shared with the IL-2 receptor and the γ_c_ subunit, which is shared with cytokines pertaining to the common γ_c_ ‘superfamily’ of cytokines, such as IL-2, IL-4 and IL-7. Both the β and γ subunits are expressed on IEL and when IL-15/IL-15Rα interacts with these receptor subunits, it elicits JAK/STAT-mediated signalling events that alter lymphocyte function in a cell type-specific manner (Perera *et al*., 2012; Mishra, Sullivan and Caligiuri, 2014). It is well established that IEL require IL-15 for their survival in the small intestine as mice lacking either IL-15 or IL-15Rα had dramatically reduced numbers of natural killer (NK) cells, NK T cells, central memory CD8+ T cells and IEL (Lodolce *et al*., 1998; Kennedy *et al*., 2000).

But IL-15 has been shown to affect more than just IEL survival. Specifically, it was reported that as compared to other common γ_c_ cytokines, IL-15 induced a marked increase in proliferation, cytolytic effector function and expression of pro-inflammatory molecule IFN-γ in human IEL *in vitro* (Ebert, 1998). Further, in transgenic mice overexpressing human IL-15 under the control of an enterocyte specific promoter T3b, IEL were massively expanded, and the intestinal epithelium displayed sings of damage and villous blunting (Ohta *et al*., 2002). These effects could be reversed using a CD122-specific blocking antibody to perturb IL-15 activity (Yokoyama *et al*., 2009). IL-15 expression is chronically elevated in various tissue-specific autoimmune disorders, such as Coeliac disease (CeD) (Abadie and Jabri, 2014). CeD is an enteropathy whereby genetically susceptible individuals have an adverse reaction to gluten, causing immune-mediated damage to the small intestine. Alongside elevated levels of IL-15, one of the hallmarks of CeD is increased numbers of IEL in the small intestine, attributed to IL-15 driven proliferative expansion of IEL (Jabri *et al*., 2000; Mention *et al*., 2003; Jabri and Abadie, 2015). It was also observed that IEL derived from patients with CeD had elevated expression of activating Natural Killer (NK) cell receptors such as NKG2D and CD94 and expanded massively in response to high levels of IL-15 (Jabri *et al*., 2000; Meresse *et al*., 2004). These reports and others suggest that regulation of receptor expression is a key mechanism by which IL-15 controls IEL activity. Thus, for IEL to utilise IL-15 as a survival stimulus but also respond to rising levels of IL-15 as a ‘danger signal’, IL-15 regulation must be tightly controlled, such that it is only upregulated during tissue damage and returned to normal levels upon resolution of the infection.

Much of these data have been derived from CeD patients and ‘healthy’ controls from oesophagus-gastro-duodenoscopies for non-CeD complaints. Due to the cell contact-dependent mechanism of IL-15 presentation, it is likely that IEL from CeD patients receive multiple additional signals from the epithelial cells in which they are in contact with *in vivo*. For example, the stress-induced NKG2D ligands; MIC-A and MIC-B which are elevated on damaged epithelial cells in CeD have been shown to activate IEL (Meresse *et al*., 2004, 2006). This could explain why in studies looking at IEL isolated from CeD patients, there is enhanced cytolysis in response to IL-15, because not only have these cells been exposed to elevated levels of IL-15, but they have also been in contact with ‘stressed’ epithelial cells, which express enhanced levels of stress-induced ligands such as MIC-A and MIC-B (Groh *et al*., 1998), and likely other surface molecules indicative of tissue damage.

While studies focussing on IEL in the context of CeD have allowed insight into the role IL-15 may be playing in pathological settings, we wanted to investigate the intrinsic effects of exposure to high levels of IL-15 on ‘normal’ murine IEL to assess whether IL-15 alone was sufficient to induce the effects described in previous studies. Using high resolution mass spectrometry, we developed a proteomic map of the global changes induced in purified TCRγδ CD8αα, TCRαβ CD8αα and TCRαβ CD8αβ IEL subpopulations as a result of 24hrs exposure to high levels of IL-15. Our data have revealed new insights into how IL-15 regulates the activation status of IEL through the upregulation of various activating and inhibitory receptors, biosynthetic and bioenergetic activation, and induction of proliferation. Importantly, the data revealed a critical role for the proto-oncogenes PIM1 and PIM2 kinases in IL-15-induced proliferation, growth and acquisition of effector function of IEL.

## RESULTS

### Proteome profiling reveals distinct features of high IL-15/Rα stimulated IEL

Elevated trans-presentation of IL-15 by IEC can activate IEL cytotoxicity and proliferation *in vivo* (Ohta *et al*., 2002; Yokoyama *et al*., 2009, 2011), however, it is not clear how IL-15 drives the activation of IEL, and whether these effects are direct. To answer these questions, we used quantitative label-free high-resolution mass spectrometry to explore how IL-15 shapes the proteome of the main murine IEL subsets; TCRαβ CD8αβ, TCRαβ CD8αα and TCRγδ CD8αα (Fig. S1A). All 3 populations were sorted to greater than 96% purity. We used IL-15 coupled to its receptor the IL15Rα (IL-15/Rα) to better mimic the trans-presentation of IL-15 by IEC, and because it is better at stimulating proliferation and survival compared to soluble IL-15 (Rubinstein *et al*., 2006; Stoklasek, Schluns and Lefrançois, 2006). IEL were treated for 24hrs with 100ng/mL IL-15/Rα (3.4nM), which we designated as ‘high levels’, based on pSTAT5 induction (Fig. S1B), high survival (Fig. 1A) and proliferation induced at this concentration. Untreated controls were derived directly *ex vivo*, as IEL do not survive well in culture in the absence of IL-15, and there was also substantial cell death at low concentrations of IL-15 (Fig. 1A) which would affect the quantification of the proteomic data. However, we have subsequently confirmed key findings both in *ex vivo* vs IL-15 stimulated samples, as well as in IEL cultured in low (2-10ng/mL) vs high levels (100ng/mL) of IL-15/Rα.

**Figure 1.**
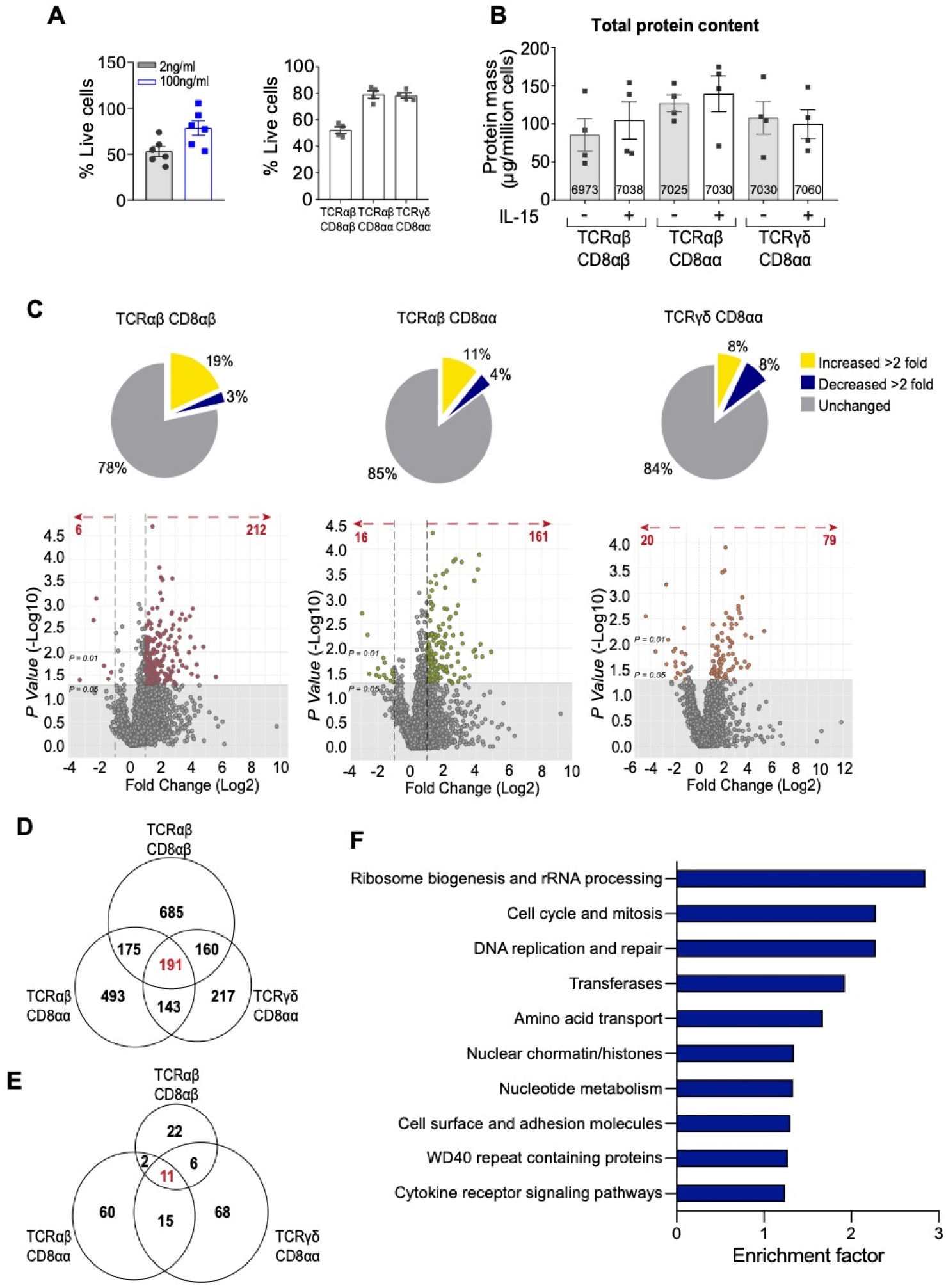
Global effects of IL-15/Rα stimulation on the proteomes of the three main IEL subsets. (**A**)Bar chart shows the percentage of live cells following 24hrs IL-15/Rα (100ng/mL) stimulation of CD8α+ IEL (left) and FACS-purified IEL subsets (right) used for mass spectrometric analyses. Percentages were calculated from the number of cells that were considered live (negative for DAPI staining) following IL-15/Rα treatment (100ng/mL or 2ng/mL) for 24hrs, divided by the number of cells seeded for culture (1 million/mL). (**B**)The protein content of each IEL subset before and after 24hrs IL-15/Rα stimulation, protein content is presented as µg/million cells. The number of proteins identified in each subset are displayed on the respective bars. (**C**) Pie charts show the fold change in expression of proteins in IL-15-treated samples compared to untreated samples for each IEL subset. Proteins with a fold change >2 were considered upregulated while proteins with a fold change <0.5 were downregulated, and those in between were deemed ‘unchanged’ by IL-15 stimulation. Volcano plots show differential expression of proteins following IL-15 stimulation for each IEL subset. Data are presented as the distribution of the copy number ratio (IL-15-treated vs untreated) (log2 (fold change)) against the inverse significance value (-log10(*p-value*)). Proteins were considered upregulated following IL-15 stimulation if the log2 fold change value was >1, and similarly proteins were considered downregulated if the log2 fold change was <-1. The grey area depicts the cut off for proteins deemed to have an insignificant fold change (p-value >0.05). Hence, proteins above the grey area were deemed significantly changed following IL-15 stimulation. (**D**) and (**E**) Venn diagrams show the commonality of proteins upregulated and downregulated (>2-fold), respectively, across all IEL subsets following 24hr IL-15/Rα stimulation. (**F**) Top 10 functional clusters enriched in proteins that were commonly 2-fold upregulated by IL-15/Rα stimulation in all 3 subsets, as in (D). See also supplementary table 2. All error bars are s.e.m, proteomic data derived from 4 biological replicates.

We identified and quantified >7100 proteins in the total data set (4 biological replicates/population; +/- IL-15), providing the first high-resolution quantitative proteomes of 3 developmentally diverse IEL populations. The intensities associated with the proteins identified in different biological replicates of each condition were reasonably well correlated; the lowest *r*^2^ value was 0.76 (Fig. S1C). For subsequent interpretation of the data, intensities were converted into estimated copy number per cell using the proteomic histone ruler method (Wiśniewski *et al*., 2014). Protein copy numbers were then used to calculate the protein content as mass (µg/million cells), and despite their differences in ontogeny, all IEL subsets had remarkably similar protein content that was not significantly changed by IL-15 (Fig. 1B). Moreover, there was also a high correlation of protein copy numbers between subsets with and without IL-15 stimulation (Fig. S1D). These data indicate that different IEL subsets are very similar in their protein make-up and overall protein abundance.

To better understand how IL-15 differentially altered the proteomic landscape of each subset, we calculated the fold change in expression of proteins (IL-15-treated vs untreated) and found that ∼15% of the total proteome of CD8αα-expressing IEL changed >2 fold in response IL-15 stimulation. More specifically, these changes were equally distributed between upregulated and downregulated proteins in TCRγδ IEL, whereas in TCRαβ CD8αα, >11% of the proteome were upregulated >2 fold. IL-15 had the largest impact on the total proteome of TCRαβ CD8αβ IEL, with ∼19% of the total proteome being upregulated >2 fold, and ∼3% downregulated >2 fold (Fig. 1C). The strong impact on TCRαβ CD8αβ IEL was surprising, as these cells expressed the lowest copy numbers of IL-2Rβ (CD122), necessary for signal transduction downstream of IL-15 (Fig. S1E). However, the phosphorylation of STAT5 both *ex vivo* and in response to IL-15/Rα stimulation was comparable in all 3 subsets (Fig. S1F). It is likely therefore that the IL-15 receptor levels on TCRαβ CD8αβ IEL are sufficient to activate downstream signalling at a similar level to that seen in the other IEL subsets.

To identify global IL-15 targets in IEL, we next asked which proteins were commonly regulated by IL-15 among IEL subsets. We identified 191 proteins that were commonly upregulated >2fold irrespective of significance, in all three IEL subsets, and >330 proteins commonly upregulated in 2 or more subsets (Fig. 1D). On the other hand, IL-15/Rα stimulation led to the downregulation of just a handful of proteins, mainly inhibitory proteins, e.g. serpins, galectins and E3 ubiquitin ligases. In addition, epithelial proteins such as Villin 1 and glycoprotein A33 were downregulated, potentially contaminants in the *ex vivo* samples that were lost upon culture (Fig. 1E). Functional annotation clustering revealed that IL-15 most commonly upregulated proteins involved in cell cycle, DNA replication and repair, ribosome biogenesis, transferases, including kinases, synthases and methyltransferases, and cell surface receptors and adhesion molecules among others (Fig. 1F and Supplementary Table 2). In summary, these data reveal for the first time many new pathways in the IL-15-regulated proteome of IEL.

### IL-15 drives IEL proliferation by triggering G1/S transition

IEL maintained in high levels of IL-15/Rα (100ng/mL) over 96hrs increased in numbers (Fig. 2A), whereas low levels (2ng/mL) appeared only enough to maintain decreasing numbers of IEL. This indicated that high levels of IL-15 either enhanced survival or proliferation, or both, of IEL. As previously established in human refractory coeliac disease (RCeD) IEL cell lines (Malamut *et al*., 2010), we also found that IEL expressed the anti-apoptotic factors Bcl2 (∼200,000 copies), Bcl-xL (5,000-10,000 copies), and Mcl1 (10,000-30,000 copies), supporting cell survival. However high IL-15/Rα did not significantly enhance their expression above what is expressed basally *ex vivo* (Fig. 2B). We then assessed proliferation using CellTrace™ CFSE and found that IEL start to divide at 48hrs in response to 100ng/mL but did not divide at all in low levels of IL-15/Rα, even after 4 days (Fig. 2C). These data support the notion that high IL-15 signals are required for inducing IEL proliferation, as suggested by the functional annotation of the proteomic data (Fig. 1F). In order to propel resting T cells through the G1 restriction point and into the S phase of the cell cycle, the cyclin-dependent kinase inhibitor 1B (CDKN1B), also known as p27 needs to be phosphorylated and degraded. Once phosphorylated and targeted for degradation, p27 is unable to bind to and inhibit downstream cell cycle targets D-type cyclins and the associated cyclin-dependent kinases 4 and 6 (CDK4/6). Further analyses revealed that IL-15-treated IEL had elevated levels of CDK4/6 (Fig. 2D). Moreover, the ratio of p27 to cyclins D2 and D3, an indicator of G1/S transition, was strongly decreased in IEL by IL-15/Rα (Fig. 2E). DNA synthesis measurements using EdU in IEL cultured in low or high IL-15/Rα confirmed that only high IL-15 levels triggered entry of IEL into the S phase of the cell cycle after 48hrs (Fig. 2F). Overall these data indicate that high IL-15 trigger proliferation of IEL by regulating key cell cycle checkpoint proteins to allow IEL to exit from quiescence.

**Figure 2.**
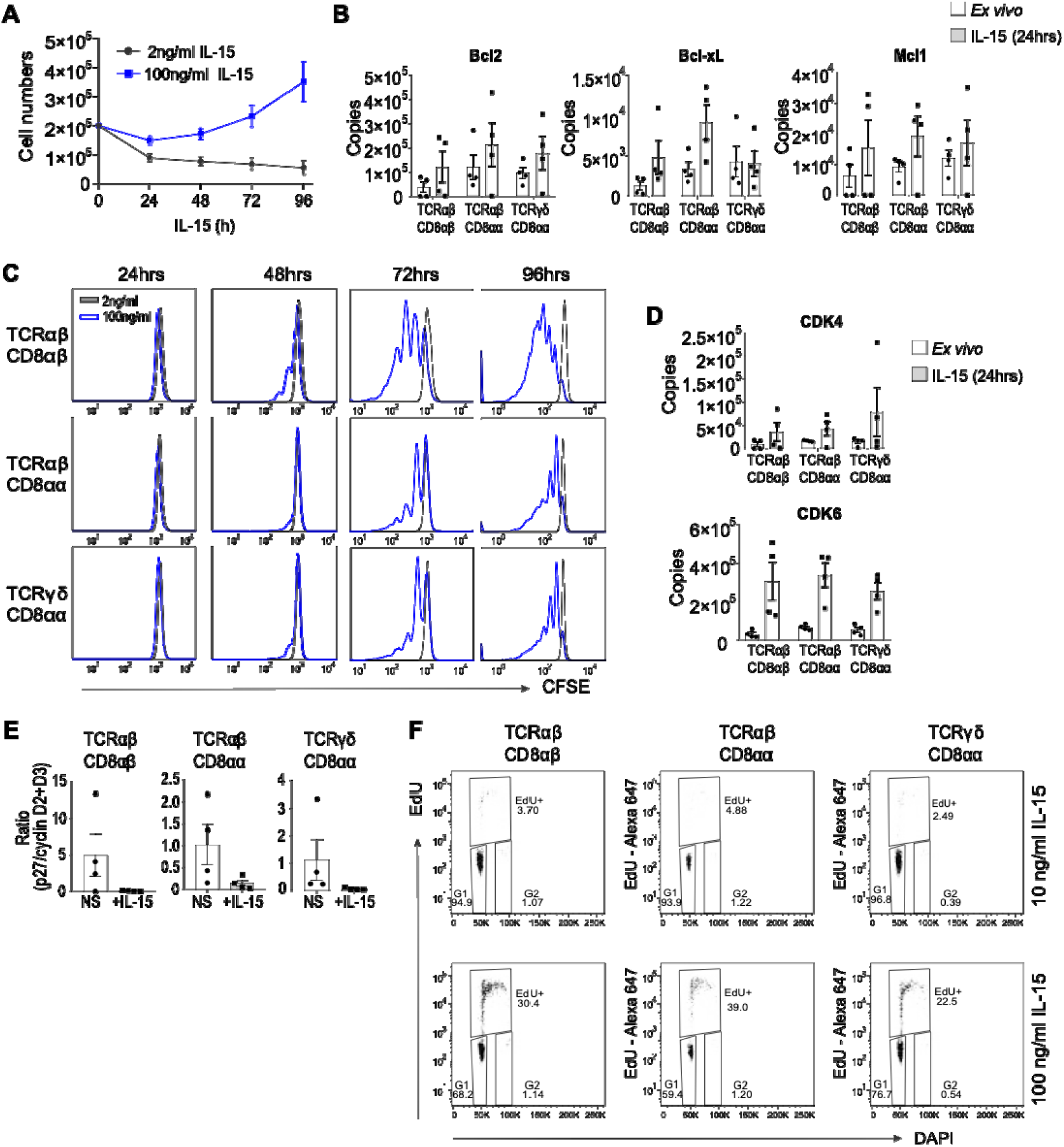
High IL-15/Rα stimulation drives proliferation of IEL by licensing the G1/S transition. (**A**) Total numbers of cells obtained on each day after culture of CD8^+^ IEL (200,000 cells (200µl/well)) with IL-15/Rα (100ng/mL or 2ng/mL). Data are the mean and s.e.m of 8 independent cultures. (**B**) Estimated copy numbers per cell of survival proteins Bcl-2, Bcl-xL and Mcl1 in each IEL subset prior to and following 24hrs 100ng/mL IL-15/Rα stimulation. (**C**) IEL were isolated, enriched for CD8α expression and stained with CellTrace™ CFSE prior to stimulation with either 2ng/mL (grey) or 100ng/mL (blue) IL-15/Rα for 4 days. Every 24hrs cells were stained for subsets; TCRαβ CD8αβ, TCRαβ CD8αα, TCRγδ CD8αα and CFSE expression was analysed by flow cytometry. The discrete peaks in the histograms represent successive generations of live, DAPI-negative IEL. (**D**) Estimated copy numbers of cyclin-dependent kinases 4 and 6 (CDK4/6) prior to and following 24hr 100ng/mL IL-15/Rα stimulation. (**E**) Bar graphs showing the ratio of p27 molecules to the total sum of cyclin D2 and cyclin D3 molecules in each IEL subset prior to and following 24hr 100ng/mL IL-15/Rα stimulation. (**F**) IEL were isolated, enriched for CD8α expression and stimulated with either 10ng/mL or 100ng/mL IL-15/Rα for 48hrs. Cells were stained for IEL subsets as in (C) and DNA synthesis was assessed by incorporation of EdU. Data is representative of 2 biological replicates. All error bars are s.e.m, proteomic data derived from 4 biological replicates, symbols on the bars represent the replicates.

### High IL-15 stimulation triggers biosynthetic pathways in IEL

A key feature of T cell activation is a massive increase in cell size fuelled by an increase in the uptake of nutrients and metabolic reprogramming (MacIver, Michalek and Rathmell, 2013). IEL already bear many hallmarks of activation, including high Granzyme expression, CD69 and CD44 expression but are still small, metabolically quiescent cells (Konjar *et al*., 2018). Although we did not detect an increase in the overall protein content (100-150 μg/million cells) of IEL at the 24hr treatment used in the proteomic experiment (Fig. 1B), IEL maintained in high levels of IL-15/Rα increased in size and granularity by 48hrs and continued to grow over 96hrs, as compared to those maintained in low levels (Fig. 3A). Therefore, we interrogated the proteomic data to gain insights into the biosynthetic and bioenergetic machinery and mechanisms by which IL-15 might drive this expansion in cell size.

**Figure 3.**
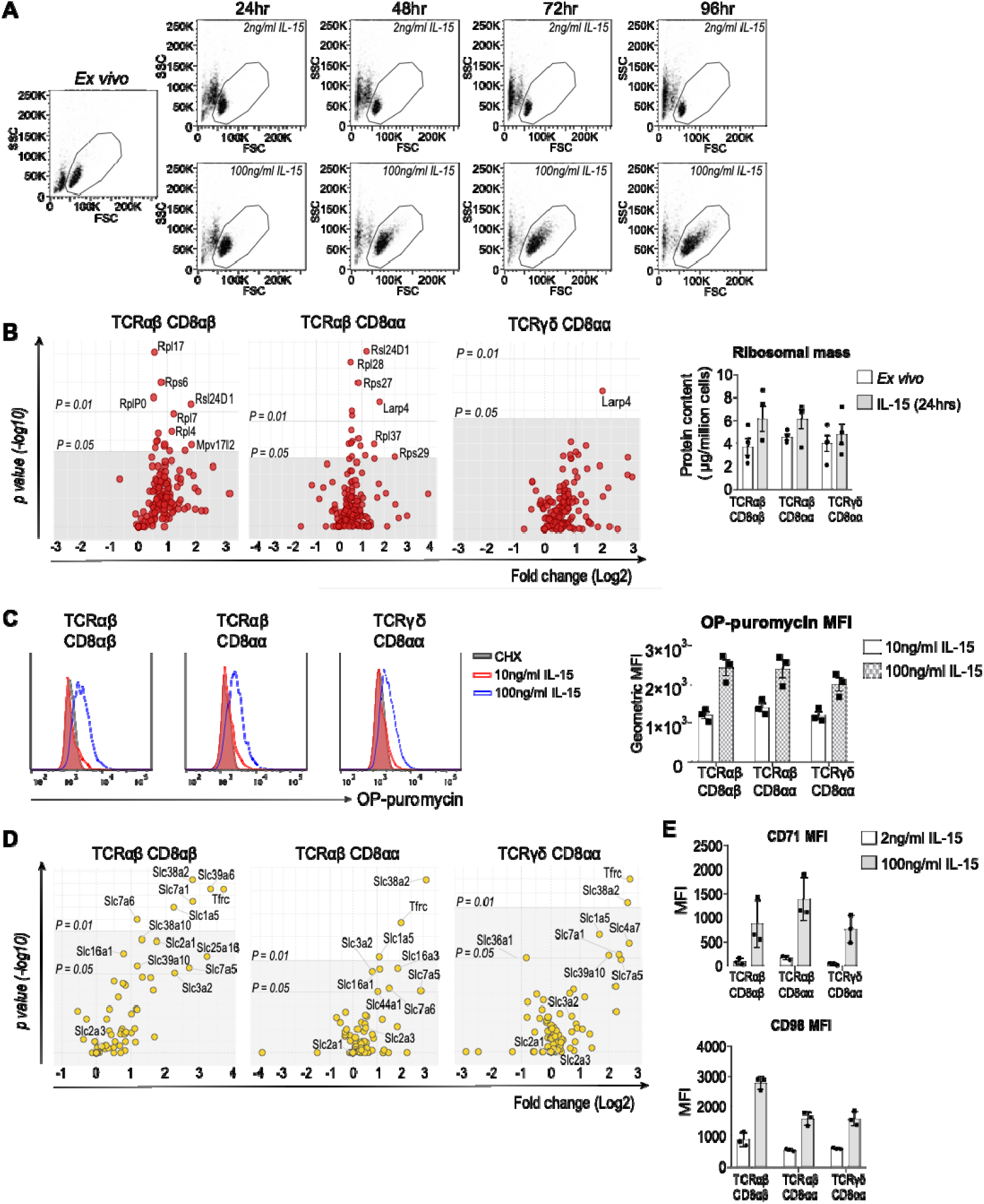
IL-15/Rα increases IEL protein synthesis, nutrient uptake and growth. (**A**) IEL were isolated, enriched for CD8α and cultured in either low levels (2ng/mL) or high levels (100ng/mL) IL-15/Rα. Dot plots show the forward scatter (FSC, indicator of cell size) vs side scatter (SSC, indicator of granularity) of live IEL every 24hrs over 96hrs culture. (**B**) Volcano plots were generated showing the differential expression of ribosomal proteins (GO:0005840) in IEL subsets. Data are presented as the distribution of the copy number ratio (IL-15-treated vs untreated) (log2 fold change) against the inverse significance value (-log10 (*p*-value)). Bar graphs show the protein content of the sum of ribosomal proteins in non-stimulated vs IL-15-treated IEL. (**C**) OPP incorporation in CD8α+ IEL cultured with low (10ng/mL) or high (100ng/mL) IL-15/Rα for 48hrs. As a negative control, incorporation was inhibited by cycloheximide (CHX) pre-treatment. OPP incorporation was assessed by flow cytometry after 15 minutes administration. Histograms show OPP incorporation in CHX-treated IEL (grey), IEL treated with low levels of IL-15/Rα (red) and IEL treated with high levels of IL-15/Rα (blue). Bar graph shows the MFI of OPP-Alexa fluor 647 in each IEL subset. Histograms are representative of 3 biological replicates. (**D**) Volcano plots were generated showing differential expression of nutrient transporters following IL-15 stimulation for each IEL subset as in (B). (**E**) Flow cytometric analyses of CD71 and CD98 expression on IEL cultured in either low levels (2ng/mL) or high levels (100ng/mL) IL-15/Rα for 72hrs. Data is 3 biological replicates. All error bars are s.e.m, proteomic data derived from 4 biological replicates, symbols on the bars represent the replicates.

The most significantly enriched gene set in our dataset corresponded to rRNA synthesis and ribosome biogenesis. Closer analyses revealed that ribosomal proteins (GO: 0005840) were largely upregulated by IL-15/Rα, with an overall increase in the total ribosome content (µg/million cells), particularly in the TCRαβ CD8αβ subset (Fig. 3B). We also noted that key ribosome biogenesis factors such as Nsa2 and Rrs1 were enriched more than 2-fold 24hrs after IL-15/Rα stimulation in all 3 subsets (Supplementary table 1). Since ribosomal biogenesis is synonymous with protein translation, we also assessed the effects of high IL-15 stimulation on protein synthesis. We applied O-propargyl-puromycin (OPP) labelling to assess global protein synthesis in single cell assay by flow cytometry. OPP is an analogue of puromycin that is incorporated into nascent polypeptide chains and can be labelled with a fluorophore, thus permitting measurement of rates of translation. Strikingly, only IEL cultured in 100ng/mL IL-15/Rα for 48hrs showed an increase in protein synthesis after 15 minutes of OPP-labelling, whereas cells cultured in low IL-15 did not show any labelling above cycloheximide (CHX) treated controls (Fig. 3C). Thus, high levels of IL-15 signalling trigger protein synthesis in IEL, that is supported by ribosome biogenesis.

Given the increase in protein synthesis in IL-15-stimulated IEL, we next assessed how IL-15 altered the expression of limiting nutrient transporters that provide biosynthetic precursors for anabolic processes. We observed a substantial increase in the expression of various nutrient transporters including the transferrin receptor (*Tfrc*, CD71), large neutral amino acid transporter, SLC7A5/LAT1, the Y+L amino acid transporter SLC7A6, and neutral amino acid transporters, SL38A2 and SLC1A5 (ASCT2), upon IL-15/Rα stimulation (Fig. 3D), many of which are key nutrient transporters for T cell activation (Carr *et al*., 2010; Sinclair *et al*., 2013; Nakaya *et al*., 2014; Bröer, Fairweather and Bröer, 2018). We confirmed these data by flow cytometry for CD71 and for CD98, the heavy chain associated with SLC7A transporters (Fig. 3E). These data suggest that IL-15 complexes enhance the ability of IEL to take up amino acids, the building blocks for protein synthesis, and other nutrients essential for optimal T cell function, such as iron and zinc (Preston *et al*., 2015; Colomar-Carando *et al*., 2019).

Conventional T cells increase their capacity for both glycolysis and oxidative phosphorylation upon activation to satisfy their increased bioenergetic and biosynthetic demands (Ma *et al*., 2019). Therefore, we asked if IL-15 also induced similar changes in IEL. IL-15 marginally induced the upregulation of glucose transporter GLUT1 from 1000-2000 copies to ∼5000-10000 copies per cell (Fig. 4A). However, this was still very low compared to what was found on activated CD8 T cells (50,000 molecules per cell) (Howden *et al*., 2019). Conversely, we noted that IEL basally expressed significantly higher levels of the high affinity glucose transporter GLUT3 (∼130,000 molecules per cell) as compared to GLUT1 (Fig. 4A), and in contrast to conventional activated T cells which have been shown to preferentially express high levels of GLUT1 (Howden *et al*., 2019). Thus, IEL constitutively have the potential to take up glucose, and this is enhanced by elevated IL-15 signalling (Fig. 4B). IEL did not upregulate any known enzymes in glucose metabolising pathways except for the glucose phosphorylating enzyme hexokinase 2 (HK2), which was expressed at low levels (8000-16000 copies per cell) (Fig. 4C) but is nevertheless thought to contribute to glycolysis in T cells (Tan *et al*., 2017). The proteomic data also revealed a small increase in the lactate exporters MCT1 and MCT3 (Fig. 4C). To investigate whether IEL metabolise glucose through glycolysis, we measured lactate output in IEL cultured in high or low levels of IL-15 and found that high levels of IL-15 specifically increased production of lactate (Fig. 4D). These data indicate that the small, but significant changes in GLUT1, HK2, and lactate transporter expression translate to activation of glycolysis in IL-15 stimulated IEL.

**Figure 4.**
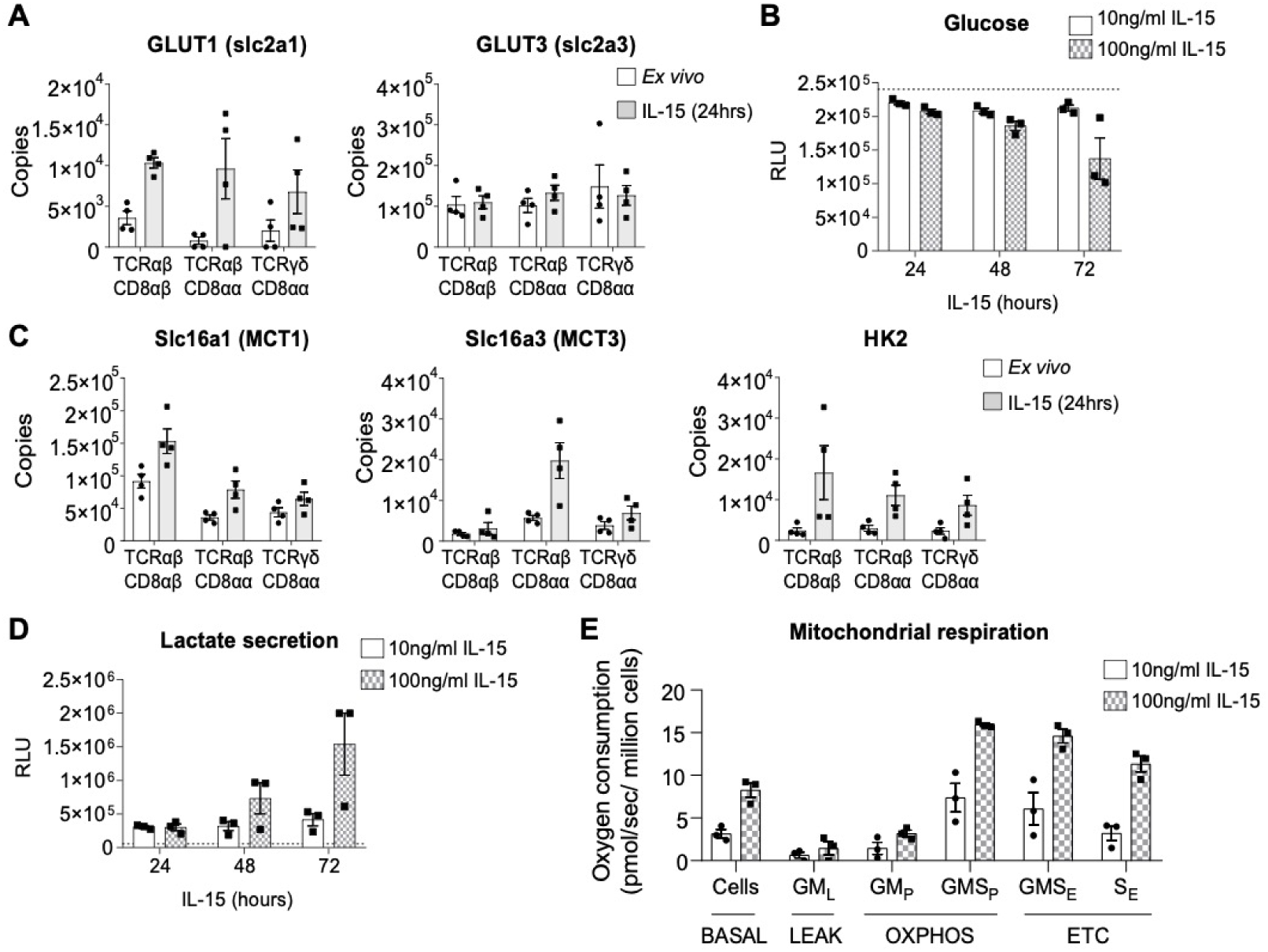
High IL-15/Rα stimulation increases mitochondrial respiration in IEL. (**A**) Estimated protein copy numbers of the glucose transporters, GLUT1 and GLUT3, for all IEL subsets either *ex vivo* (white bars) or 24hr IL-15/Rα stimulated (grey bars). (**B**) IEL were cultured in low (10ng/mL) or high (100ng/mL) IL-15/Rα for 72hrs. Media was collected every 24hrs and levels of glucose remaining in the medium were assessed by bioluminescence assay. The dotted line depicts the signal from the control wells containing only medium but no cells. (**C**) Estimated protein copy numbers of the lactate transporters Slc16a1 (MCT1) and Slc16a3 (MCT3) and the rate limiting glycolytic enzyme hexokinase 2 (HK2). Data is shown for all IEL subsets either *ex vivo* (white bars) or 24hr IL-15-stimulated (grey bars). (**D**) Lactate output from IEL cultured as in (B). (**E**) Mitochondrial respiration measurements in IEL cultured in low (10ng/mL) or high (100ng/mL) IL-15/Rα for 44hrs. Oxygen consumption is expressed as pmol/ (sec x million cells). Respiratory rates were measured in cells (BASAL), then the cells were permeabilized with 10μg/mL Digitonin, and mitochondrial respiratory rates measured after the subsequent addition of glutamate and malate (GM_L_), ADP to stimulate respiration (GM_P_) along with succinate to stimulate complex II (GMS_P_), uncoupler (FCCP) to measure maximal electron transport (GMS_E_), rotenone to block complex I (S_E_) and Antimycin A to inhibit complex III. The residual oxygen consumption after rotenone and Antimycin A treatment was subtracted from all values shown here. OXPHOS, oxidative phosphorylation, ETC, electron transfer capacity. All error bars are s.e.m, symbols on the bars represent replicates.

Interestingly, IL-15 also increased expression of several proteins involved in mitochondrial respiration, including components of the mitochondrial ribosome and electron transport chain (Supplementary table 2). We therefore assessed mitochondrial respiratory capacity of IL-15/Rα-stimulated IEL using high-resolution oxygraphy. The data show that high IL-15 triggers an increase in basal oxygen consumption at the cellular level (Fig. 4E, BASAL). Further analyses were performed in permeabilised cells to permit addition of exogenous mitochondrial complex substrates and ADP. This approach allowed us to evaluate the contributions of individual mitochondrial complexes to the increased mitochondrial respiration. Based on these analyses, we could attribute the increase in mitochondrial respiration to an increase in the coupling of complex II to complexes III and IV, and also to increased electron transfer capacity (ETC). These data indicate that IL-15 stimulated increased mitochondrial function, which therefore resulted in increased spare respiratory capacity of IL-15 stimulated IEL, and an overall increase in oxidative phosphorylation (OXPHOS). Taken together, high IL-15 triggered multiple pathways to enable the switch from quiescent to active IEL.

### Cytotoxic effector function of IL-15/Rα-stimulated IEL

Thus far we have focussed on the enriched pathways in the Il-15 regulated IEL proteome. However, a key question in IEL biology is how IL-15 triggers cytotoxicity of IEL towards surrounding epithelial cells to contribute to the pathology of CeD. Previous studies have indicated that the cytotoxic potential of human IEL is enhanced by exposure to high levels of soluble IL-15 (Ebert, 1998; Roberts *et al*., 2001). The enhancement of IEL cytotoxicity by IL-15 has been attributed to an increase in GzmB expression in IEL, and to changes in NK receptor expression. Our proteomic data revealed that IEL expressed very high levels of GzmA and GzmB in the resting state (12-50 million and 7-10 million copies per cell respectively) and levels of perforin that were comparable to effector T cells ((Hukelmann *et al*., 2016) and Fig. 5A). IL-15 led to a modest upregulation of GzmB expression, however it is unclear what functional relevance this increase would have on IEL effector potential given that they already express such high amounts of Granzymes under normal conditions. We confirmed the modest increase in GzmB by flow cytometry, but also detected a decrease in GzmA protein in IEL cultured for 24hrs in high IL-15/Rα (Fig. 5B). We further investigated the expression of several key proteins involved in degranulation, including the Munc family of proteins, Stxbp2 (MUNC18-2), and Unc13d (MUNC13-4), and the Rab GTPase Rab27a. None of these proteins were regulated by IL-15 (Fig. 5B), confirming the idea that IEL are already fully primed for activation and degranulation.

**Figure 5.**
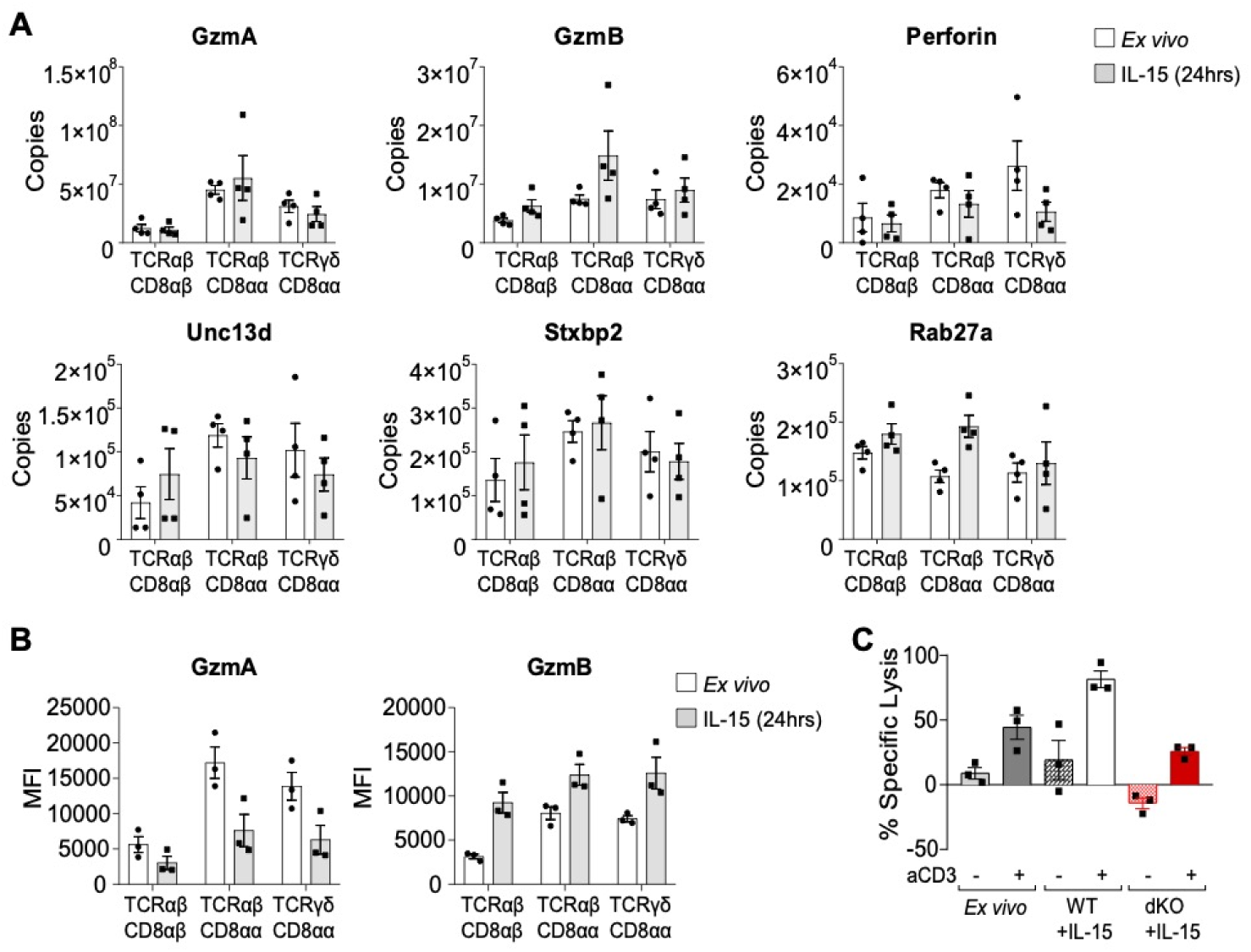
IL-15 effects on IEL cytotoxicity. (**A**) Bar graphs show estimated protein copy numbers of cytotoxic molecules GzmA, GzmB, Perforin, Unc13d, Stxbp2 and Rab27a. Data is shown for all IEL subsets either *ex vivo* (white bars) or 24hr IL-15-stimulated (grey bars). (**B**) Intracellular expression of GzmA and GzmB was analysed by flow cytometry in freshly isolated IEL compared to those cultured with 100ng/mL IL-15/Rα for 24hrs. All error bars are s.e.m, proteomic data derived from 4 biological replicates, symbols on the bars represent the replicates. (**C**) Luciferase-transduced K562 cells were co-cultured for 24hrs with freshly isolated IEL or WT and GzmA/B dKO IEL that had been pre-treated with 100ng/mL IL-15/Rα (+/- aCD3) for 72hrs, at an effector to target (E:T) ratio of 40:1. Bar graphs represent the percentage of specific lysis for each condition. Error bars are s.e.m, and data is 3 biological replicates.

Given these data, we asked whether IL-15 also enhanced cytotoxicity in murine IEL, and whether this was dependent on Granzymes. WT IEL *ex vivo* were not efficient killers in a redirected killing assay against K562 target cells in the absence of TCR stimulation via anti-CD3 treatment (Fig. 5C, E:T ratio of 40:1). Culturing IEL in 100ng/mL IL-15/Rα for 72hrs enhanced IEL ability to kill, however maximal killing was dependent on additional stimulation via the TCR. To investigate whether IEL utilise Granzymes to kill, we performed cytotoxic killing assays using GzmA^-/-^GzmB^-/-^ (GzmA/B dKO) mice. In contrast to WT cells, GzmA/B-dKO IEL treated with high levels of IL-15 failed to kill any of the target cells and even had the opposite effect of increasing K562 cell viability. This effect was only poorly reversed by TCR stimulation (Fig. 5C). These results suggest that GzmA/B are crucial for IEL killing and that other cytotoxicity-inducing molecules such as TRAIL and FasL do not replace granzymes in this context. Taken together, these data indicate that IL-15 acts as a costimulator for TCR-induced cytotoxicity that is mediated by granzymes, however it also suggests that granzyme upregulation is unlikely to be the mechanism involved.

### Il-15 upregulates multiple activating and inhibitory receptors on IEL

Previous studies have reported that IEL derived from patients with CeD had elevated expression of activating Natural Killer (NK) cell receptors NKG2D and CD94 (Jabri *et al*., 2000; Meresse *et al*., 2004). Interestingly, we did not detect an increase in the protein copy number of CD94 on any IEL subset and only observed a modest increase of NKG2D exclusively in TCRαβ CD8αα IEL (Fig. 6A, B). Using flow cytometry, we confirmed that NKG2D was not expressed by IEL in the resting state, but expression was upregulated in a small percentage (∼2-8%) of TCRαβ CD8αα IEL specifically following IL-15/Rα treatment, and this did not increase drastically even after 72hrs of culture (Fig. 6A). Similarly, CD94 was expressed mainly by ∼5-15% of TCRαβ CD8αα and TCRγδ CD8αα after IL-15 stimulation (Fig. 6B). Thus, CD94 and NKG2D are unlikely to be direct targets of IL-15 in the murine small intestine.

**Figure 6.**
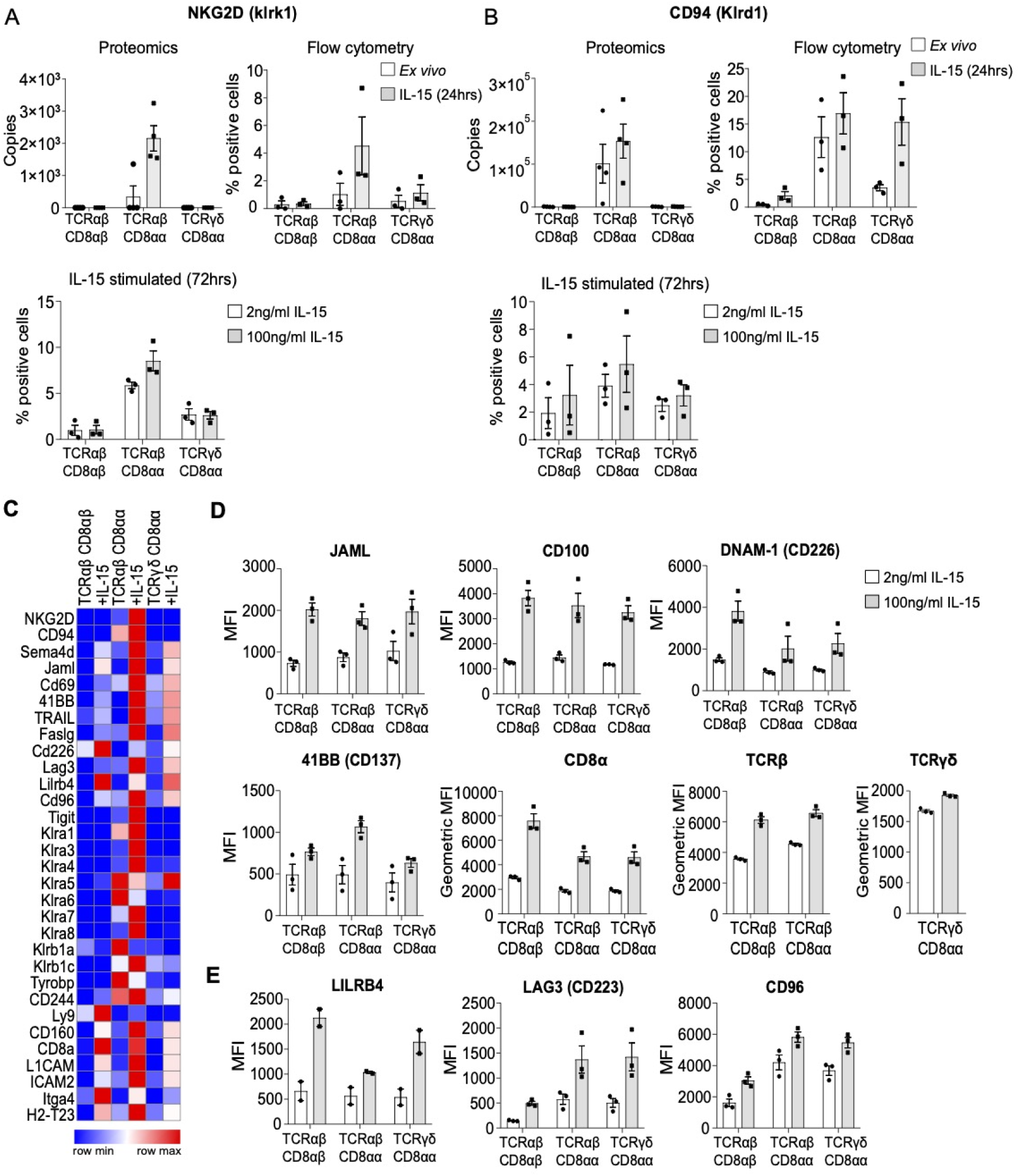
Activating/inhibitory receptor expression following high IL-15/Rα stimulation. Estimated protein copy numbers of (**A**) NKG2D and (**B**) CD94, with corresponding flow cytometric analysis on either *ex vivo* vs 24hr IL-15-stimulated IEL or IEL that had been cultured in low levels (2ng/mL) or high levels (100ng/mL) IL-15/Rα for 72hrs. Flow cytometry data are presented as % positive cells following gating on IEL subsets; TCRαβ CD8αβ, TCRαβ CD8αα, TCRγδ CD8αα. (**C**) A list of surface receptors identified in the proteomics was compiled and estimated copy numbers per cell for each biological replicate were averaged. These data were used to generate a heat map showing the expression profile of these surface receptors across IEL subsets. (**D**) CD8α+ IEL were cultured in 2ng/mL or 100ng/mL IL-15/Rα for 72hrs and assessed for their expression or various surface receptors identified in the heatmap. Flow cytometric data are shown as MFI for (**D**) activating receptors; JAML, CD100, DNAM-1, 41BB, CD8α, TCRβ and TCRγδ and (**E**) inhibitory receptors; LILRB4, LAG3 and CD96. All error bars are s.e.m, proteomic data derived from 4 biological replicates, flow cytometry data derived from 3 biological replicates, with the exception of LILRB4 of which 2 biological replicates are presented.

IEL are known to express a range of other surfaces receptors and the proteomic data revealed that many of these were significantly upregulated following IL-15 stimulation in all IEL subsets (Fig. 6C). Significantly, a number of these receptors have the potential to impact upon the activation and cytotoxic effector function of IEL such as adhesion molecules (e.g. ICAM2, Integrin α4, L1CAM) and activating receptors (e.g. JAML, CD100, 4-1BB and CD226). For example, the junctional adhesion molecule-like (JAML) receptor was significantly upregulated following IL-15 stimulation in all subsets and has been reported to act as a co-stimulatory receptor on TCRγδ T cells in the skin, potentially by recruiting PI3K (Verdino *et al*., 2010, 2011; Witherden *et al*., 2010). To investigate JAML expression at the single cell level among each IEL subset, we used flow cytometry and found that not only was JAML uniformly upregulated on all IEL in each subset following 24hr IL-15 stimulation compared to *ex vivo* IEL, this increased expression was sustained when comparing IEL cultured in high vs low IL-15/Rα for 72hrs (Fig. 6D). We also detected upregulation of other activating receptors such as CD100, 4-1BB and CD226 (DNAM-1) (Fig. 6D). CD226 is particularly interesting in the context of cytotoxic function, as its ligation is thought to trigger cytotoxic activation of NK cells. Moreover, both TCR and CD8α expression were increased quite substantially by IL-15, both in the proteomics (Supplementary Table 1) and by flow cytometry at 72hrs (Fig. 6D). On the other hand, we also noted that high IL-15 stimulation led to the upregulation of various inhibitory receptors such as LAG3, LILRB4 and CD96 (Fig. 6C, E). Thus, the potential of IL-15 to activate IEL *in vivo* may be driven through increased adhesion to epithelial cells, and the balance of both activating and inhibitory signals received through multiple receptors besides NKG2D.

### Identification of PIM1/2 kinases as regulators of IEL responses to elevated IL-15 signals

Among proteins that were not expressed in IEL *ex vivo* but upregulated substantially by high IL-15/Rα were the PIM kinases, PIM1 and PIM2 (Fig. 7A). Using immunoblotting, we confirmed a clear induction of all 3 isoforms of PIM1, and 2 isoforms of PIM2 in IEL stimulated with high IL-15/Rα (Fig. 7B). PIM proteins are serine/threonine kinases transcriptionally regulated downstream of receptors activating pathways such as the JAK/STAT signalling pathway (Basham *et al*., 2008). While PIMs have been shown to play a key role in proliferation, cell survival and protein synthesis, their functions are highly cell type-specific and a role in IEL has not yet been defined. PIM1^-/-^/PIM2^-/Y^ (PIM1/2 dKO) mice are healthy and have normal complement of lymphocytes in the spleen (Fig. S2). To assess the contributions of PIM kinases to IL-15-stimulated IEL, we isolated IEL from PIM1/2 dKO mice. As IL-15 is crucial for IEL survival, it was possible that IL-15 might regulate IEL survival via PIM kinases, given that the PIM kinases have been shown to play a role in cell survival via apoptosis inhibition (Macdonald *et al*., 2006). However, IEL numbers and composition were normal in PIM1/2 dKO mice (Fig. 7C), and their survival was unperturbed in response to low levels of IL-15 (Fig. 7D). On the other hand, responses to high levels of IL-15 were severely impaired in PIM1/2 dKO mice. The numbers of WT IEL increased over 96hrs in response to high but not low levels of IL-15, however PIM1/2 dKO IEL failed to increase in numbers comparatively (Fig. 7E). We used CellTrace™ CFSE reagent to track proliferation and found that while all 3 IEL subsets from WT mice began dividing by 48hrs, this proliferation was completely abolished in PIM1/2 dKO mice (Fig. 7F). This defect was specific to IEL, as activated splenic T cells proliferated normally in the presence of IL-2 (Fig. S2B-D), another γc cytokine that also induces the PIM kinases (Rollings *et al*., 2018). Additionally, we noted that loss of either PIM1 or PIM2 on their own did not have as strong an effect on IEL proliferation, although the PIM1 KO IEL were clearly still delayed in their proliferative response to high IL-15/Rα (Fig. S3A, B).

**Figure 7.**
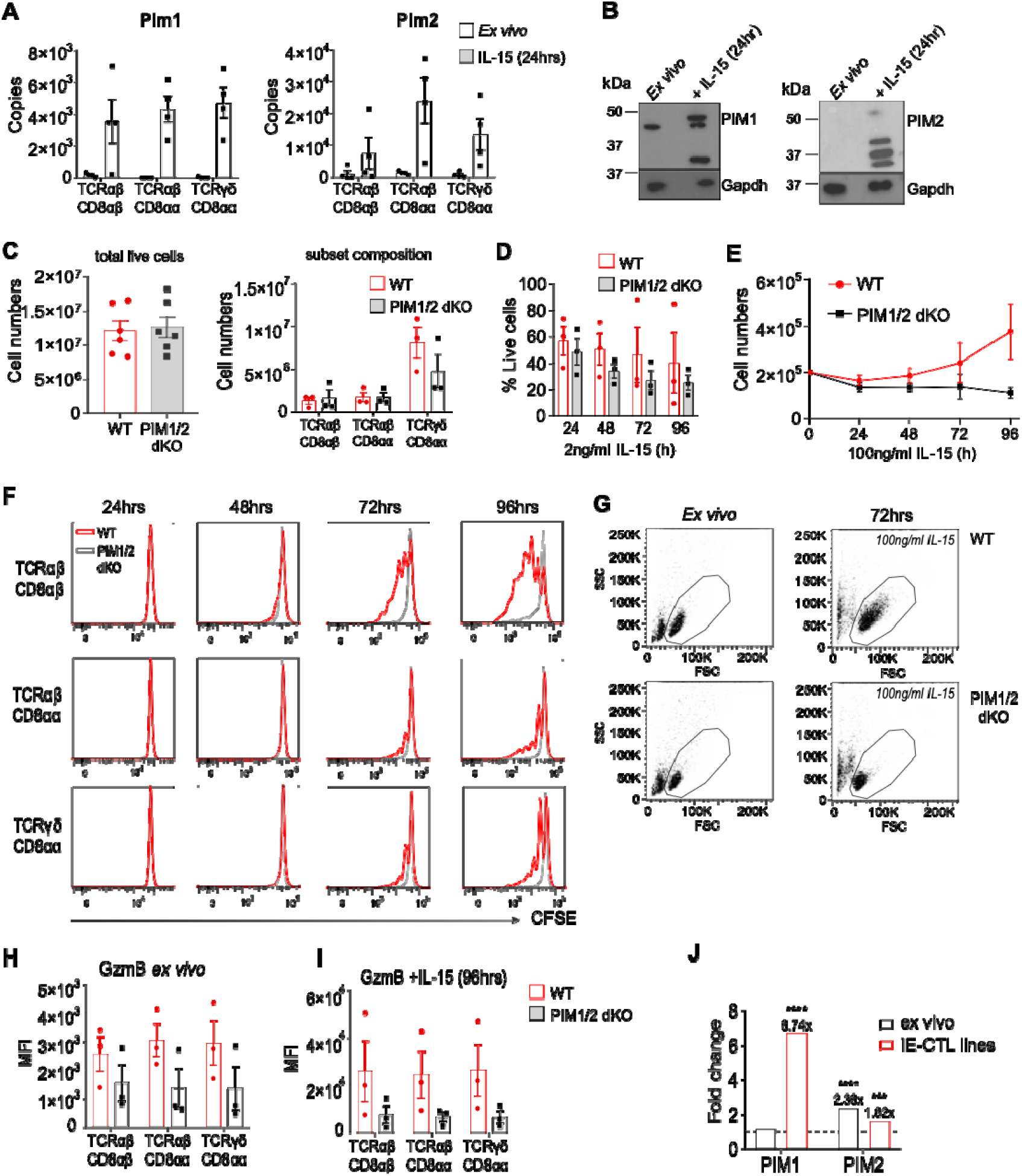
PIM kinases regulate IEL responses to high IL-15/Rα stimulation. (**A**) Estimated copy numbers per cell are shown for the PIM1 and PIM2 kinases in IEL subsets that were either untreated (*ex vivo*) or maintained in 100ng/mL IL-15/Rα for 24hrs. (**B**) Western blot data showing PIM1 (right) and PIM2 (left) expression in 20µg of *ex vivo* IEL or 24hr IL-15-stimulated IEL. Antibodies against GAPDH were used as a loading control, data is representative of 3 experiments (**C**) Bar graphs show the absolute cell counts (left; 6 biological replicates) and subset composition (right; 3 biological replicates) of IEL isolated from WT (red) and PIM1^-/-^/PIM2^-/Y^ (PIM1/2 dKO) (grey) mice. (**D**) Bar graph shows the percentage of live IEL from either WT or PIM1/2 dKO mice that were cultured in low (2ng/mL) IL-15/Rα for 0-96hrs. Percentages were calculated from the number of cells that were considered live (negative for DAPI staining) following IL-15/Rα treatment divided by the number of cells seeded for culture (1 million/mL) at each timepoint. Data is 3 biological replicates. (**E**) Line graph shows the cell numbers of IEL from either WT or PIM1/2 dKO mice that were cultured in high (100ng/mL) IL-15/Rα for 0-96hrs, data is 4 biological replicates. (**F**) IEL were isolated from WT and PIM1/2 dKO mice, enriched for CD8α expression and stained with CellTrace™ CFSE prior to stimulation with 100ng/mL IL-15/Rα for 4 days. Every 24hrs cells were stained for subsets and CFSE expression was analysed by flow cytometry. The discrete peaks in the histograms represent successive generations of live, DAPI-negative IEL. (**G**) Dot plots show the forward scatter (FSC) vs side scatter (SSC) of live IEL *ex vivo* as compared to IEL cultured in 100ng/mL IL-15/Rα for 72hrs from both WT (top) and PIM1/2 dKO (bottom) mice. (**H**) *Ex vivo* and (**I**) IEL that were cultured in 100ng/mL IL-15/Rα from both WT and PIM1/2 dKO mice were stained for intracellular GzmB expression, presented as MFI. (**J**) CD8+ intraepithelial T-cell lines or *ex vivo* single-cell suspensions from the epithelial compartment isolated from human duodenal biopsies were stimulated with human recombinant IL-15 for 2 hours and RNA sequencing performed. Bar graphs show the fold change in gene expression (untreated vs IL-15-treated) for PIM kinases; PIM1 and PIM2. Data derived from GSE120904 and Ciszewski et al, 2019. PIM1/2 dKO experiments are 3 biological replicates unless otherwise stated. Proteomic data are 4 biological replicates, points on bar graphs represent the biological replicates. Error bars are s.e.m.

We also noted that IEL lacking PIM1/2 failed to grow in size and granularity as compared to WT IEL exposed to high levels of IL-15 (Fig. 7G). As previously shown, IEL express extremely high levels of GzmA and B in the resting state, however, PIM dKO IEL had a significant reduction in the levels of GzmB (Fig. 7H), but not GzmA (Fig. S2E). Granzyme levels increase in WT IEL that have been exposed to IL-15 in long-term cultures, however PIM1/2-deficient IEL failed to upregulate GzmB even in the presence of high levels of IL-15 for 96hrs (Fig. 7I). These data indicate that the PIM kinases may regulate growth and protein synthesis in IEL, which may contribute to PIM kinase regulation of GzmB expression in IEL.

Finally, we asked whether human IEL also upregulate PIM kinase expression in the context of IL-15 and/or CeD. Analyses of a previously published publicly available RNA-seq datasets on human TCRαβ CD8αβ IEL cell lines, and on *ex vivo* IEL stimulated with high levels of soluble IL-15 indicated that human IEL also upregulated PIM1 and PIM2 upon IL-15 stimulation, at least at the mRNA level, although the level of expression varied somewhat between the *ex vivo* IEL and the cell lines ((Ciszewski *et al*., 2020), and Fig. 7J). Furthermore, data mining of an earlier published study of gene expression profiling of human IEL and peripheral blood lymphocytes (PBL) from CeD patients revealed that both PIM1 and PIM2 are expressed at higher levels in IEL compared to PBL ((Meresse *et al*., 2006), and Fig. S2F). Together these data indicate that upregulation of PIM kinases by IL-15 may be a conserved pathway for regulating IL-15 responses in IEL.

## DISCUSSION

Of all T lymphocytes, IEL are the least understood, with limited insights into their biology and regulation. Using high-resolution mass spectrometry, we here reveal the IL-15-regulated proteomes of three distinct IEL subsets; TCRαβ CD8αβ, TCRαβ CD8αα and TCRγδ CD8αα. By focussing on IL-15, a key cytokine that activates IEL function, we dissect the pathways and modules regulating IEL activation, proliferation and proteome remodelling. We observed that the major effect of IL-15 complexes on IEL is to trigger their proliferation, not only by inducing the cell cycle machinery, but also by activating all of the accompanying changes required, such as protein and nucleotide synthesis, ribosome biogenesis, and targeted bioenergetic activation to fuel these changes. IL-15 induced cell surface proteins potentially involved in epithelial interactions, particularly cell adhesion molecules, and cytotoxic activators. Importantly, our work identified the PIM1 and PIM2 kinases as key regulators of IEL function. We show for the first time that proliferation and growth of IEL are dependent on the PIM1 and PIM2 kinases, something that does not appear to hold true for conventional T cells.

IEL are unusual T cells that have an ‘activated’ T cell phenotype, in that they express activation markers such as CD69 and they harbour huge amounts of cytotoxic molecules. Despite these hallmarks, IEL do not exhibit effector function in the resting state. With no known endogenous antigenic stimuli, it has been difficult to define the mechanisms necessary for IEL activation. IL-15 has been shown induce cytolytic effector function in human IEL in the context of certain target cells (Ebert, 1998; Roberts *et al*., 2001). It has previously been suggested that IL-15 triggers IEL cytotoxic activity by upregulating expression of Granzyme B, and the activating receptor NKG2D, both in human and murine IEL (Meresse *et al*., 2006, Abadie *et al*., 2020). However, in our proteomics data, we find only modest upregulation of NKG2D, on a very small proportion of IEL. We also reveal that Granzyme B is expressed at approximately 10 million molecules per cell prior to IL-15 stimulation, so it is not clear how upregulating expression of Granzyme B further is necessary for activating IEL cytotoxicity. Admittedly there are differences between human and mouse IEL. For example, NKG2D is expressed on resting human IEL, whereas it is not expressed in murine IEL. However, we also find that IL-15 increases the cytolytic activity of murine IEL dramatically, particularly if the TCR is also engaged. Moreover, mice overexpressing IL-15 in the intestine display signs of villous atrophy, suggesting that murine IEL can be activated to kill epithelial cells in the presence of elevated levels of IL-15, yet even in these settings, less than 10%of IEL were shown to express NKG2D (Ohta *et al*., 2002; Yokoyama *et al*., 2009). Therefore, other as yet undiscovered pathways must be contributing to IL-15 driven cytotoxicity of IEL.

Our data revealed upregulation of various activating receptors, such as JAML and CD100 in response to 24hr IL-15 stimulation. JAML was recently reported to act as a costimulatory receptor for epidermal TCRγδ T cells, increasing proliferation and cytokine production upon interaction with its ligand CAR on keratinocytes (Witherden *et al*., 2010). Similarly, CD100 was shown to specifically regulate intestinal TCRγδ IEL activation in the context of colitis and epidermal wound repair (Witherden *et al*., 2012; Meehan *et al*., 2014). CD226 is particularly striking as it is also expressed on human IEL, and its activation on NK cells and CD8 T cells has been variously described to trigger cytotoxicity, proliferation, and cytokine production (Shibuya *et al*., 1996; Zhang *et al*., 2015). In our proteomic data, these receptors were not only expressed by TCRγδ IEL but shared by all IEL subsets and IL-15 stimulation commonly increased their expression, highlighting potential mechanisms for IL-15-induced activation of IEL. Another possible explanation for the increased cytotoxic activity of IL-15 stimulated IEL is that IL-15 lowers the signalling threshold required for TCR or other co-receptor activation. In this context, it is interesting to note that only culture in high IL-15 levels upregulated TCR expression in IEL. The involvement of the TCR and the various co-stimulatory and inhibitory receptors on IEL in triggering IEL cytotoxicity remains to be resolved.

One fundamental insight from our proteomic data was that IL-15 induced the upregulation of PIM1 and 2 in response to IL-15. Using PIM1/2-deficient mice we have shown that IEL depend on both PIM kinases for their proliferative expansion and growth in response to IL-15 *in vitro*. Interestingly, PIM1/2-deficient IEL appeared phenotypically normal in unchallenged mice, except for a slight reduction in GzmB, but not GzmA, expression. This fits with the data that we did not detect much PIM1 or PIM2 expression in *ex vivo* IEL, despite the basal level of IL-15 signalling that must be ongoing *in vivo*. It appears that there is a threshold of IL-15 signalling required to trigger the expression of PIM kinases. It is also noteworthy that in the absence of PIM kinases, IL-15 could not induce upregulation of GzmB expression. These data suggest that the PIM kinases also control the cytotoxic activation of IEL.

Intraepithelial lymphocytosis and villous atrophy, both hallmarks of CeD, are thought to be driven by IL-15. Currently the only treatment available for CeD patients is a life-long gluten-free diet. However, issues due to cost, non-compliance with the diet, and gluten contamination cause symptoms in up to 30-50% of CeD patients. Additionally, in a small percentage of CeD patients, even being gluten-free does not reduce inflammation, leading to refractory coeliac disease (RCeD) (Meresse, Malamut and Cerf-Bensussan, 2012). In RCeD, overexpression of IL-15 persists and abnormal IEL accumulate, causing extensive damage to the intestinal epithelium and a poor prognosis. An estimated 40% of RCeD patients go on to develop enteropathy associated T cell lymphomagenesis, where the lymphomas are IEL-derived. Hence, there is an urgent need for treatments for CeD, RCeD, and associated lymphomas. Currently, IL-15 blocking antibodies are being investigated as a treatment strategy (Vicari *et al*., 2017), however, with limited success (Cellier *et al*., 2019; Lähdeaho *et al*., 2019), thus identification of new targets for small molecule inhibitors would be beneficial. Our data suggest that inhibitors against the PIM kinases may also be suitable targets for the treatment of CeD and RCeD. PIM kinase inhibitors are currently in clinical trials for the treatment of certain types of hematological malignancies and solid cancers (Cortes *et al*., 2018), thus there is clear potential for these drugs to also be used for the treatment of CeD and its complications.

This study provides the first comprehensive proteomic analyses of the effects of high levels of IL-15 on IEL. The effects of IL-15 on IEL receptor expression and biosynthetic pathways, among others, are novel pathways that warrant further investigation. Moreover, the identification of the PIM kinases for IEL biology in particular is an unanticipated finding of this study. Further research will reveal how PIM kinases orchestrate IEL biology and immune responses, and whether this a specific feature of only these lymphocytes. The proteomics data and insights obtained in this study will serve as an important resource, not only for the study of IEL in CeD, but also in the context of infection, inflammatory bowel diseases, and cancer where IL-15 upregulation has been seen.

## Acknowledgments

This study was supported by a Wellcome Trust PhD studentship to OJJ (215309/Z/19/Z). MV and MS are supported by the Wellcome Trust and Royal Society (Sir Henry Dale Fellowship to MS, 206246/Z/17/Z). JMM is supported by Marie Sklodowska-Curie (705984) and EMBO (ALTF 1543-2015) fellowships. We would like to thank A. Rennie, A. Gardner and R. Clarke for cell sorting and D. Campbell, R. Gourlay and J. Varghese for mass spectrometric analyses in the MRC-PPU.

## METHODS

### Mice

C57BL/6 mice were purchased from Charles Rivers and acclimatised for a minimum of 10 days prior to use in experiments. PIM1^-/-^ /PIM2^-/Y^ (PIM1/2 dKO), PIM1^-/-^ (PIM1 sKO) and PIM2^-/Y^ (PIM2 sKO) mice were obtained from Cantrell Lab; University of Dundee, Scotland. GzmA^-/-^/GzmB^-/-^ (GzmA/B dKO) were obtained from Julian Pardo; University of Zaragoza, Spain. All mice were bred and maintained under specific pathogen-free conditions in the Wellcome Trust Biocentre at the University of Dundee in compliance with U.K. Home Office Animals (Scientific Procedures) Act 1986 guidelines.

### Isolation of IEL from the intestinal epithelium

IEL were isolated as described in (James, Vandereyken and Swamy, 2020). Briefly, small intestines were dissected from proximal duodenum to terminal ileum and using a gavage needle, flushed with 20mL cold PBS to remove luminal contents. Intestines were cut longitudinally and then transversely into small 5-10mm pieces. The pieces were placed into 25mL complete RPMI media (RPMI +10% FBS, 1% Pen/Strep & L-glutamine) with 1mM DL-Dithiothretiol (DTT) and incubated on a shaker for 40 minutes at RT. After centrifugation and vortexing in complete RPMI media the cells were passed through a 100µm strainer and isolated cells were centrifuged in a 36%/67% Percoll/PBS density gradient at 700g for 30 minutes with no brake. The IEL appear as diffuse layer at the interface between the two Percoll concentrations.

### Sample preparation for mass spectrometry

For proteomics experiments, 4 biological replicates were generated and IEL were isolated from 8 mice per biological replicate (male, C57BL/6, aged 10-12 weeks). IEL were isolated from the small intestine as previously described. For further enrichment of the CD8α^+^ IEL population, following Percoll density gradient centrifugation, an EasySep™ Release PE positive selection kit (STEMCELL Technologies) was used with a PE-conjugated anti-mouse CD8α antibody (BioLegend) as per the manufacturer’s instructions. Cells were stained with PerCP eFluor 710 (TCRγδ), APC (TCRβ), PE (CD8α), FITC (CD8β) and PE-Cy7 (CD4) for isolation of pure populations of TCRγδ CD8αα, TCRαβ CD8αα and TCRαβ CD8αβ IEL using fluorescent activated cell sorting (FACS) (Supplementary Fig. 1A). We purified ∼10million TCRγδ CD8αα, ∼5million TCRαβ CD8αα and ∼5 million TCRαβ CD8αβ per biological replicate. 2 million cells from each purified cell population were pelleted and snap frozen in liquid N_2_ and the remaining cells treated with 100ng/mL IL-15/Rα for 24hrs as previously described then similarly pelleted and snap frozen.

IEL cell pellets (both IL-15-treated and untreated) were lysed in 200µl lysis buffer (4% SDS, 10mM TCEP, 50mM TEAB (pH 8.5)). Lysates were boiled and sonicated (15 cycles of 30s on/30s off) and protein concentrations determined by EZQ^®^ Protein Quantitation Kit (Invitrogen). Lysates were then alkylated with iodoacetamide (IAA) for 1hr at room temperature in the dark. For protein clean-up, 200µg SP3 beads (Hughes *et al*., 2014) were added to lysates before elution in digest buffer (0.1% SDS, 50mM TEAB (pH 8.5), 1mM CaCl_2_) and digested with LysC and Trypsin, each at a 1:100 (µg enzyme:protein) ratio. Peptide clean-up was performed according to the SP3 protocol. Samples were resuspended in 2% DMSO and 5% formic acid. For fractionation, off-line high pH (9.5) reverse phase chromatography was used. Using an Ultimate 3000 HPLC (Thermo Scientific), peptides were separated into 16 concatenated fractions, then manually pooled to 8 orthogonal fractions, dried and eluted into 5% formic acid for analysis by LC-MS. Samples were sent to MRC-PPU Mass Spectrometry facility, University of Dundee, where each fraction was analysed by label-free quantification (LFQ) using an LTQ OrbiTrap Velos Pro (Thermo Scientific) with a 240-minute gradient per fraction.

### Processing and analysis of proteomics data

The MS data files were processed with MaxQuant version 1.6.0.3 as described in (Rollings *et al*., 2018), against mouse reviewed proteome from Uniprot, downloaded on 25/07/2017. The data set was filtered to remove proteins categorised as “contaminants”, “reverse” and “only identified by site”. MaxQuant-derived data was converted into estimated copy numbers per cell in Perseus 1.6.1.1 as described (Wiśniewski *et al*., 2014).

### Cell culture of IEL and cytotoxicity assay

Isolated IEL were further enriched using an EasySep™ Mouse CD8α positive selection kit (STEMCELL technologies) as per the manufacturer’s instructions. IEL positively enriched for CD8 expression were resuspended in culture medium (RPMI+ 10% FBS + 1% Pen/Strep, L-glutamine, sodium pyruvate, non-essential amino acids, 2.5% HEPES and 0.1mM β-mercaptoethanol) with various concentrations of Mouse IL-15/IL-15R Complex Recombinant Protein (eBioscience™). These cells were seeded in a round-bottom 96-well plate at 1 million cells/mL (2×10^5^ cells per well) and incubated at 37°C, 10% CO_2_. For analysis of proliferation, CellTrace™ CFSE Cell Proliferation Kit (Invitrogen) was used; briefly, cells were treated with 5μM CFSE at 37°C for 10 minutes before quenching in ice cold PBS +15% FCS and put into culture as described above. For the bioluminescence-based cytotoxicity assay, luciferase-expressing K562 cells (kind gift of Dr. S. Minguet, Freiburg) were plated at a concentration of 5×10^3^ cells per well in a 96-well flat bottom plate in triplicates. 75μg/mL D-firefly luciferin potassium salt (Biosynth) was added to the K562 cells and bioluminescence was measured using a PHERAstar plate reader to establish the bioluminescence baseline. For a maximal cell death (positive control), triton X-100 was added to K562 cells at a final concentration of 1%. For spontaneous death (negative control), culture medium was added to K562 cells. For the test, WT or GzmA/B dKO IEL that had been cultured for 72hrs in 100ng/mL IL-15/Rα were added to K562 cells at a 40:1 effector-to-target (E:T) ratio and incubated for 24hrs at 37°C, 10% CO_2_. Bioluminescence was measured as relative light units (RLU). Percentage specific lysis was calculated with the following formula: % specific lysis = 100 x (average spontaneous death RLU – test RLU) / (average spontaneous death RLU – average maximal death RLU).

### Cell culture and treatments for splenocytes and lymph nodes

Single cell suspensions from WT and Pim1/2 dKO spleens were activated with anti-CD3 + anti-CD28 [both 0.5 µg/mL, clones 2C11 and 37.51 respectively, (eBioscience/Thermo Fisher Scientific)] in the presence of recombinant human IL-2 [20 ng/mL (Proleukin, Novartis)]. After 48 hrs cells were washed out of activation media, then split daily into fresh media and IL-2 (20 ng/mL) to a density of 0.3 million cells/mL. CD8+ T cell number of WT and Pim1/2 dKO T cells was counted on a FACSVerse daily from day 2 of the culture onwards. Lymph node single cell suspensions from WT and Pim1/2 dKO were labelled with 5 µM CellTrace™ Violet Cell Proliferation Kit [(CTV) Invitrogen] for 20 minutes at 37°C, washed then activated with anti-CD3 + anti-CD28 [both 0.5 µg/mL, clones 2C11 and 37.51 respectively (eBioscience/Thermo Fisher Scientific)] at 100,000 cells per well in 96-well flat-bottomed plate in 200µL total volume for 2 days.

### Measurement of extracellular metabolites in cell culture

IEL were isolated and enriched for CD8α as previously described before plating at 200,000 cells/well in a round bottom 96-well plate in 200µl medium (glucose-free RPMI to which 2mM glucose, 2mM L-glutamine 10% dialysed FBS, 1% Pen/Strep, sodium pyruvate, non-essential amino acids, 2.5% HEPES and 0.1mM β-mercaptoethanol were added). Cells were cultured at 37°C, 10% CO_2_ and every 24hrs 10µl of medium was removed, diluted in 190ul of PBS and further diluted 2.5x for glucose measurements. Samples were frozen and stored at −20°C. At the end of the experiment, samples were thawed and 50µl aliquots transferred to a 96-well assay plate (Corning Cat.#3903). Each sample was plated in duplicate for each metabolite. The metabolites were then detected using the Lactate-Glo™ and Glucose-Glo™ assays (Promega) as per manufacturer’s instructions. Luminescence was recorded using a PHERAstar plate reader and measured as relative light units (RLU).

### Flow cytometry

Cells were plated at 2×10^5^ cells per well to a 96-well dish for staining. FC block was added to each well for 5 minutes before cells were incubated with monoclonal antibodies (mAb) against cell surface markers for 15 minutes at 4°C. Cells were stained with the following antibodies specific for murine: CD45 [clone 30.F11 (BioLegend)], TCRβ [clone H57-597 (BioLegend)], TCRγδ [clone GL3 (BioLegend or eBioscience)], CD4 [clone RM4-5 (BioLegend)], CD8α [clone 53-6.7 (BioLegend)], CD8β [clone H35-17.2 (eBioscience)], CD122 (IL-15Rβ) [clone TM-b1 (eBioscience)], NKG2D [clone CX5 (BioLegend)], CD94 [clone 18d3 (BioLegend)], JAML [clone HL4E10 (BioLegend)], CD100 [clone BMA-12 (BioLegend)], CD223 (LAG3) [clone eBioC9B7W (eBioscience)] and CD85k (LILRB4) [clone H1.1 (BioLegend)], CD226 (DNAM-1) [clone 10E5 (eBioscience)], CD69 [clone H1.2F3 (eBioscience)], CD96 [clone 3.3 (BioLegend)], CD71 [clone RI7217 (BioLegend)], CD98 [clone RL388 (BioLegend)], CD178 (Fas Ligand) [clone MFL3 (eBioscience)], CD253 (TRAIL) [clone N2B2 (eBioscience)]. For fixation before intracellular staining, cells were treated with 2% PFA at 37°C for 10 minutes. For intracellular staining, cells were washed in permeabilization buffer (eBioscience) before 1hr incubation at room temperature with mAbs specific for murine: GzmB [clone GB12 (eBioscience)] or GzmA [clone GzA-3G8.5 (eBioscience)]. For detection of phospho-STAT5 by flow cytometry, cells were fixed with 2% PFA prior to any surface stains, permeabilised with 90% ice cold methanol and incubated with phospho-STAT5 (Tyr694) [clone C11C5 (Cell Signaling Technology)] for 30 minutes at room temperature (1:200 dilution). Cells were then incubated with an anti-rabbit DyLight™649-conjugated donkey secondary Ab [clone Poly4064 (BioLegend)] for 30 minutes at room temperature (1:500 dilution). To measure DNA synthesis or protein synthesis, cells were treated with 10µM baseclick 5-Ethynyl-deoxyuridine (5-EdU) (Sigma) for 2hrs or 20µM O-propargyl-puromycin (OPP) (JenaBioscience) for 15 minutes, respectively. For OPP assays, a negative control was pre-treated with 0.1mg/mL cycloheximide (CHX) for 30 minutes before adding OPP. Cells were then harvested (∼1 million cells per condition), fixed with 4% paraformaldehyde (PFA) and permeabilised with 0.5 % triton X-100 before undergoing a copper catalysed click chemistry reaction with Alexa 647-azide (Sigma). Cells were then stained with surface markers as described above and resuspended in FACS buffer (PBS + 1% FBS (+15µg/mL DAPI for cell cycle analysis)) and analysed by flow cytometry to determine the degree of incorporation of EdU or OPP. All data was acquired on a FACSVerse flow cytometer with FACSuite software (BD Biosciences) or a FACS LSR Fortessa flow cytometer with DIVA software (BD Biosciences). Data were analysed using FlowJo software (TreeStar).

### Fluorescent cell barcoding (FCB)

IEL were isolated and enriched for CD8+ as described above. Cells were resuspended at a concentration of 1million cells/mL and 500μl plated in a 24 flat bottom well plate. Cells were warmed at 37°C for 30 min before stimulated with different concentrations of IL-15/Rα for 3hrs at 37°C. After stimulation cells were directly fixed with 500μl PFA 4% 10 min at 37°C before permeabilisation with 90% ice cold methanol. During methanol permeabilisation, each sample was stained with a mix of various concentrations of amine-reactive fluorescent dyes for 40 min, on ice before quenching with PBS + 0.5%BSA (v:v). Pacific blue dye was used at a concentration of 0μg/ml, 11.1 μg/ml or 100μg/ml and DyLight800 dye at a concentration of 0μg/ml or 25μg/ml. Barcoded samples were then pooled and stained for intracellular p-STAT5 and surface markers as described in the “flow cytometry sections”. Data were acquired using CytoFlex flow cytometer and analysed using FlowJo Software. Data were analysed using the “forward deconvolution method” described in (Krutzik *et al*., 2011). Briefly, samples were differentiated based on the fluorescence intensities of each dye and then individual samples were analysed for p-STAT5 expression.

### High resolution respirometry

Mitochondrial respiration was studied in digitonin-permeabilised IELs that enables keeping mitochondria in their architectural environment. CD8+ IEL were isolated and cultured as described above, in 10ng/mL or 100ng/mL IL-15/IL-15R complexes for 42hrs. The analysis was performed in a thermostatic oxygraphic chamber at 37°C with continuous stirring (Oxygraph-2k, Oroboros instruments, Innsbruck, Austria). Approximately 1 million cells were placed in Mir05 respiration medium (0.5 mM EGTA, 3 mM MgCl_2_, 60 mM lactobionic acid, 20 mM taurine, 10 mM KH_2_PO_4_, 20 mM HEPES, 110 mM D-Sucrose, and 1 g/L of fatty acid free Bovine Serum Albumin [BSA]; pH=7.1) in the oxygraphic chamber. Respiration protocol was adapted from the Substrate-uncoupler-inhibitor titration protocol number 11 (SUIT-011) (Votion *et al*., 2012). Briefly, after the determination of the ROUTINE oxygen consumption in absence of any substrate, cells were permeabilized with digitonin (10 µg/10^6^ cells). Substrates and inhibitors were then added sequentially to determine respiratory rates. LEAK respiration was first measured by adding glutamate (10 mM) and malate (2 mM) to the chamber (GM_L_). OXPHOS respiration was then determined by first adding 2.5 mM ADP (GM_P_). The OXPHOS GM_P_ state records electron flow from the type N-pathway (NADH-generating substrates to complex I) to Q-junction that feeds electrons to complexes III and IV. The maximal OXPHOS respiration rate GMS_P_ was then measured by adding 25 mM succinate, with both type N-pathway to Q and type S-pathway (succinate, substrate of complex II) to Q being stimulated. Electron transfer system capacity (ETS) was then assayed by adding incremental doses of Carbonyl cyanide m-chlorophenyl hydrazine (CCCP) (0.05 µM steps, GMS_E_). Complex I was blocked with 0.5 µM rotenone to determine the Succinate-pathway control state (S_E_). Finally, residual oxygen consumption (ROX) was determined in presence of 2.5 µM Antimycin A. ROX was subtracted from oxygen flux as a baseline for all respiratory states. Respiratory rates were expressed as pmol O_2_ x s^-1^ x million cells^-1^.

### Immunoblots

IEL were isolated as previously described. *Ex vivo* samples were cultured with 10ng/mL Mouse IL-15/IL-15R Complex Recombinant Protein (eBioscience™) (IL-15/Rα) for 2hrs to allow for a basal level of PIM expression before harvesting 5 million cells and pelleting. Stimulated IEL were cultured as described in “cell culture and treatments for IEL” section with 100ng/mL of IL-15/Rα for 24hrs and then similarly refreshed in medium + IL-15/Rα for 2hrs before harvesting. Cell pellets were lysed at between 50-60 million cells/mL RIPA lysis buffer (50mM Tris pH 7.4, 1% (v/v) NP-40, 0.5% (w/v) Na deoxycholate, 0.1% (w/v) SDS, 150mM NaCl, 2mM EDTA, 50mM NaF, 1mM TCEP, 5mM Na-pyrophosphate, 10mM Na-β-phosphoglycerate, 1mM Na Orthovanadate, cOmplete™ mini EDTA-free protease inhibitor tablet (Roche)) for 10 minutes on ice before centrifuging for 12 minutes at 4°C. Protein concentrations were determined using the Coomassie (Bradford) Protein Assay Kit (ThermoFisher) as directed. Lysates were boiled with NuPAGE® LDS sample buffer (4X) for 3 minutes at 100°C. 20μg of lysates were separated using a 4% stacking/10% resolving SDS-PAGE gel at 120V for 100 minutes. Following separation, proteins were transferred to a PVDF membrane for 90 minutes at 300mA. Membranes were then blocked in 5% (w/v) milk (Marvel)/PBS-Tween for 30 minutes at RT. Following blocking, membranes were washed and incubated with primary monoclonal antibodies; anti-mouse PIM1 [clone 12H8 (Santa Cruz Biotechnology)] or anti-mouse PIM2 [clone 1D12 (Santa Cruz Biotechnology)] both used at 1:100 in 5% (w/v) milk (Marvel)/PBS-Tween. Monoclonal anti-rabbit GAPDH [clone 14C10 (Cell Signalling Technology)] was used as a loading control (1:1000 in 3% BSA/PBS Tween). Membranes were washed out of primary and incubated with HRP-conjugated secondary anti-mouse or anti-rabbit IgG (Cell Signalling Technology) at 1:5000 in 5% (w/v) milk (Marvel)/PBS-Tween, or 1:2000 in 5% BSA/PBS-Tween, respectively. Membranes were exposed to ECL (Clarity Western ECL substrate or Clarity Max™ Western ECL Substrate; BioRad) and signals were detected using a ChemiDoc imaging system.

## Data analysis and statistics

All T-tests were two-tailed and assumed unequal variance of the populations. Heat maps were generated using the Morpheus tool from the Broad Institute (https://software.broadinstitute.org/morpheus).

