## Supplementary data for "Comprehensive proteomics analyses identify PIM kinases as key regulators of IL-15 driven activation of intestinal intraepithelial lymphocytes"

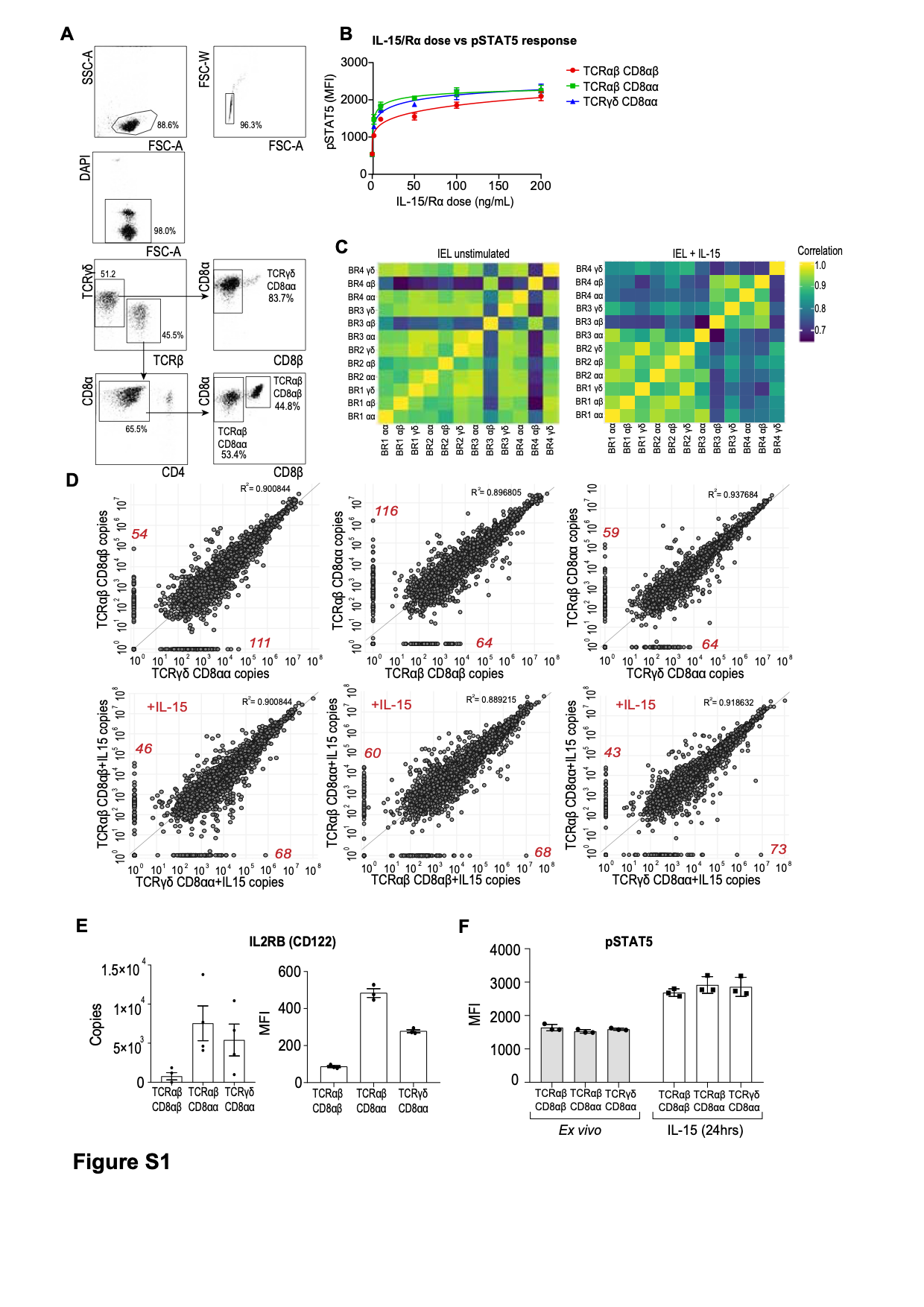
**Figure S1. Gating strategy to identify IEL subpopulations for cell sorting and global analyses of IEL proteomes**

(**A**) The gating strategy used to identify and isolate IEL subpopulations using fluorescence activated cell sorting (FACS). Cells were first gated on by size using forward scatter (FSC) vs side scatter (SSC) to separate out cells of interest from debris/non-lymphoid cells. FSC Width was used to separate out single cells from doublets. DAPI-negative cells were considered live and were further separated into three distinct IEL subpopulations based on cell surface marker expression of T cell-associated receptors; TCRγδ, TCRβ, CD4, CD8α and CD8β. The populations sorted were as follows; those that were TCRγδ+ CD8αα+, those that were both TCRαβ+ CD4- *and* either CD8αα+ or CD8αβ+. (**B**) Line graph shows an IL-15/Rα dose vs phospho-STAT5 response curve in IEL that were stimulated for 3hrs with increasing concentrations of IL-15/Rα (0ng/mL, 2ng/mL, 10ng/mL, 50ng/mL, 100ng/mL and 200ng/mL). Cells were collected and pSTAT5 levels were assessed by Fluorescent Cell Barcoding technique (FCB). Data is 3 biological replicates, and line graph is fitted with non-linear regression analysis. (**C**) Heat maps shows the Pearson’s correlation coefficient of the raw intensities of all proteins identified in the data set for each IEL subset relative to each biological replicate. A heat map was generated for both the *ex vivo* samples (top) and the IL-15-treated samples (bottom) distinctively. BR = biological replicate (e.g. BR1 = Biological replicate 1). αβ = TCRαβ CD8αβ. αα = TCRαβ CD8αα. γδ = TCRγδ CD8αα. (**D**) Protein intensities were converted into estimated copy numbers per cell using the proteomic-ruler approach and subsequently averaged across biological replicates for each IEL subset. This data was used to generate scatterplots showing the correlation between copy numbers of proteins from each of the three IEL subsets R^2^ = Coefficient for determination. (**E**) Bar charts show estimated copy numbers per cell of the β-chain of the IL-15R (CD122) in each IEL subset and flow cytometric analysis of CD122 (**F**) Flow cytometric analysis of phospho-STAT5 (Tyr694) from *ex vivo* IEL and IEL treated with 100ng/mL IL-15/Rα for 24hrs. Data presented as MFI, 3 biological replicates. Proteomic data are 4 biological replicates, points on bar graphs represent the biological replicates. Error bars are s.e.m.

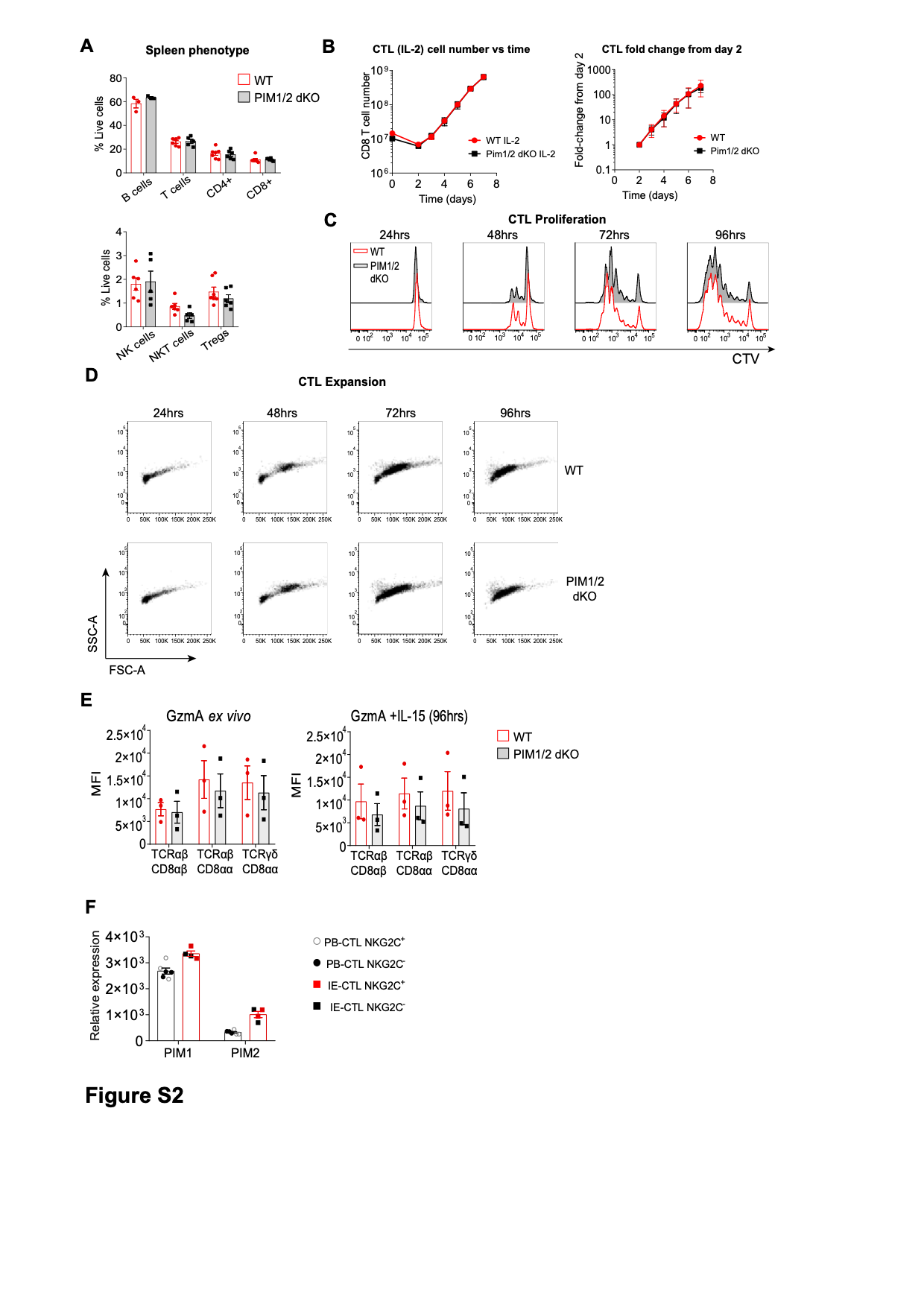

**Figure S2. Analyses of splenocytes and IL-2-stimulated CTL from PIM1/2 dKO**

(**A**) Percentage of live cells from WT and PIM1/2 dKO spleens that were B cells (B220+TCRb-), T cells (TCRb+NK1.1-) CD4 T cells (TCRb+NK1.1-CD4+), CD8 T cells (TCRb+NK1.1-CD8+), NK cells (TCRb-NK1.1+), NKT cells (NK1.1+TCRb+) or Tregs (TCRb+NK1.1-CD4+CD25+). Symbols represent biological replicates. (**B**) CD8+ T cell number of WT and PIM1/2 dKO T cells was counted on a FACSVerse daily from day 2 of culture onwards (see methods for culture conditions). Line graph on the right is cell number vs time (corrected for splitting). Data shows Mean ± SD from technical triplicates, representative of 4 independent experiments. Line graph on the left shows the fold-change in cell number from day 2 vs time Data shows Mean ± SD from 7 biological replicates, measured across 4 independent experiments. (**C**) Activated lymph node single cell suspensions from WT and Pim1/2 dKO were labelled with cell trace violet (CTV) and cell proliferation (CTV dilution) analysed by flow cytometry daily. (**D**) Corresponding cell size and granularity was measured by FSC-A and SSC-A, respectively. Data shows representative plots from 2 independent experiments. (**E**) IEL from both WT and PIM1/2 dKO mice were isolated and stained for intracellular GzmA expression *ex vivo* (left) and following 96hrs in culture with 100ng/mL IL-15/Rα (right). Bar graphs show the MFI of GzmA. Data is 3 biological replicates. (**F**) Gene expression profiling by microarray of *Pim1* and *Pim2* genes obtained from Gene Expression Omnibus dataset GSE4592. Data show microarray analysis of NKG2C^+^ and NKG2C^−^ intraepithelial cytotoxic T cells (IE-CTL) and peripheral blood cytotoxic T cells (PB-CTL) clones generated from four celiac patients.

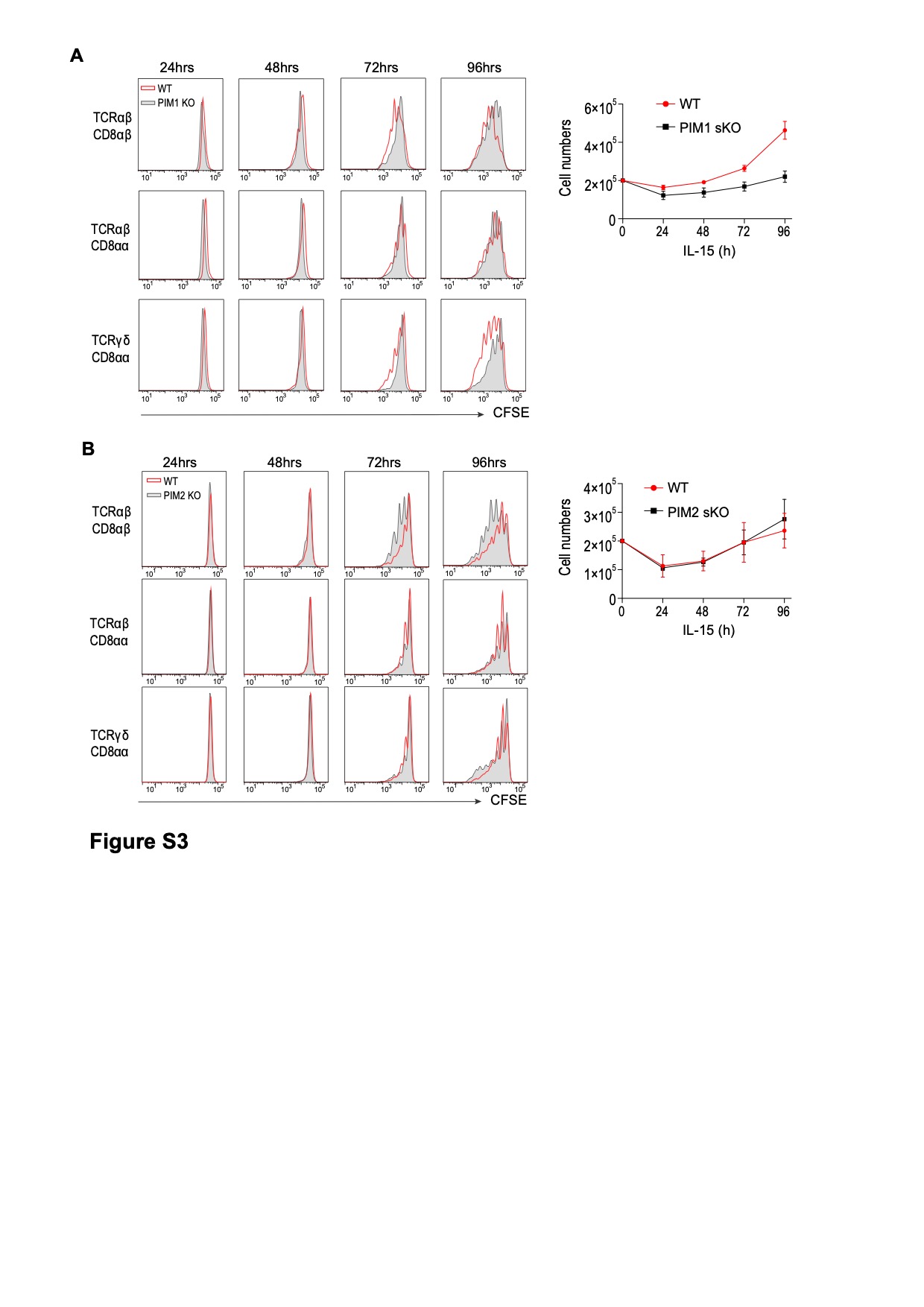
**Figure S3. IEL from PIM1^-/-^ or PIM2^-/-^ single KO (sKO) proliferate**

IEL were isolated, enriched for CD8α expression and stained with CellTrace™ CFSE prior to stimulation with 100ng/mL IL-15/Rα for 4 days. Every 24hrs cells were stained for subsets; TCRαβ CD8αβ, TCRαβ CD8αα, TCRγδ CD8αα and CFSE expression was analysed by flow cytometry. The discrete peaks in the histograms represent successive generations of live, DAPI-negative IEL for (**A**) PIM1 sKO and (**B**) PIM2 sKO IEL. Line graphs show the survival of IEL from either (**A**) PIM1 sKO, or (**B**) PIM2 sKO mice as compared to IEL from wildtype mice that were cultured in high (100ng/mL) IL-15/Rα for 0-96hrs.

**Supplementary table 1. Table with all proteins changed at least 2-fold by IL-15/Rα stimulation in each of the three main IEL subsets**

**Supplementary table 2. Functional annotation clustering of proteins upregulated in all 3 subsets at least 2-fold by IL-15/Rα stimulation**
